# Sensory Change Detection Unfolds Within a Shared Low-Dimensional Brain-State Landscape

**DOI:** 10.64898/2026.11.18.767177

**Authors:** Hadi Choubdar Parvin, Kimford J Meador, Babak Razavi

## Abstract

Detecting behaviorally relevant sensory changes requires the coordination of modality-specific sensory processing with shared attentional and control systems, yet how these processes are organized and evolve across large-scale brain networks remains unclear. Here, we investigated the connectivity architecture and temporal dynamics of auditory and visual target detection using fMRI acquired during matched oddball tasks. Cross-modal partial least-squares analysis revealed a low-dimensional architecture comprising two dominant, modality-general dimensions and additional modality-biased components. The shared dimensions distinguished a sensory–attention–action configuration from a higher-order associative and contextual configuration, whereas modality-biased dimensions showed distinct organization involving frontoparietal and attentional networks. Projecting these connectivity patterns onto time-resolved BOLD activity revealed a common dominant latent space with graded regions of preferential occupancy across modalities. Overlap-adjusted event-related analyses further showed that auditory and visual targets were associated with a similar initial displacement along the first temporal latent dimension, followed by modality-dependent trajectories, particularly along the second dimension. This later divergence reflected different balances of regional contributions within a largely shared cortical architecture. By integrating dimensionality reduction of high-dimensional functional connectivity with network and temporal analyses, our findings link the spatial organization of distributed cortical systems to their evolving responses, revealing how shared brain networks can support both common and modality-dependent aspects of sensory target detection.

## 1. Introduction

Rapid detection of behaviourally relevant changes in the sensory environment is essential for adaptive attention and action. This process is commonly examined using oddball paradigms, in which infrequent target events occur within a stream of repeated standard stimuli^1^. Although early processing depends on modality-specific sensory systems, auditory and visual targets also recruit a distributed set of temporoparietal, inferior frontal, insular, cingulate, parietal and premotor regions^2,3^. This overlap has been interpreted as evidence for a supramodal system that links sensory change detection to attentional orienting and response preparation. Consistent with this account, large-scale models of attention distinguish ventral systems involved in detecting and reorienting toward salient events from dorsal and frontoparietal systems that maintain goal-directed selection and task control^4,5^.

However, cross-modal overlap does not imply that auditory and visual changes are represented identically. Modality-specific sensory responses remain prominent, and both visual and auditory change detection engage interactions between sensory cortices and higher-order frontal regions^6,7^. Recent cross-modal studies further suggest a hierarchical organization in which brief or simple deviations retain stronger sensory-specific contributions, whereas more complex or behaviourally relevant targets recruit increasingly sustained supramodal responses^8–10^. It therefore remains unclear whether auditory and visual change detection recruit the same fixed supramodal network, converge from separate sensory pathways onto common attentional hubs, or occupy a shared architecture whose dominant dimensions are modality-general while secondary dimensions preserve sensory-specific context.

Resolving this question requires moving beyond regional activation overlap. Conventional target-versus-standard contrasts identify where responses occur but provide limited information about how distributed regions are coordinated, whether shared and modality-sensitive processes form coherent connectivity patterns, or whether these patterns evolve during ongoing task performance. Task-based functional connectivity studies indicate that cognitive operations emerge through structured modifications of large-scale network architecture^11^. In parallel, dynamic connectivity and latent-state approaches suggest that whole-brain activity evolves through structured, low-dimensional state spaces rather than as a sequence of independent regional responses^12–14^ [Song et al., 2023]. Whether auditory and visual change detection unfold within the same latent dynamical space remains unknown. Moreover, a rare target could either induce entry into a distinct target-specific state or transiently perturb an ongoing shared trajectory before the system returns toward its dominant operating regime^15,16^.

Here we asked whether auditory and visual change detection are supported by distinct modality-specific systems or organized within a shared low-dimensional architecture that preserves sensory context. We projected cross-modal functional connectivity patterns from fMRI into a common latent subspace to identify modality-general and modality-biased dimensions, then examined the spatial organization of these dimensions through their network connectivity. We next tracked their temporal expression to determine how auditory and visual task states evolve within the same dynamical space. This framework allowed us to test whether sensory modality partitions change detection into separate network configurations or are expressed as structured variation within a shared architecture, and whether target events induce distinct states or transiently perturb ongoing trajectories.

## 2. Results

### 2.1. Functional connectivity patterns in auditory and visual modalities

Seventeen healthy participants completed auditory and visual change detection tasks during fMRI scanning. Each participant performed three runs per condition; however, one auditory run was excluded due to a technical issue during acquisition. This resulted in a total of 50 auditory and 51 visual runs being included in the final analysis (Fig. 1A). Functional data were parcellated using a symmetric 180-region version of the Glasser atlas^17^. For each run, a functional connectivity (FC) matrix was computed by correlating the mean time series across all pairs of regions. These matrices were then averaged across runs within each condition to generate mean FC matrices for the auditory and visual tasks (Fig. 1B).

**Fig. 1.**
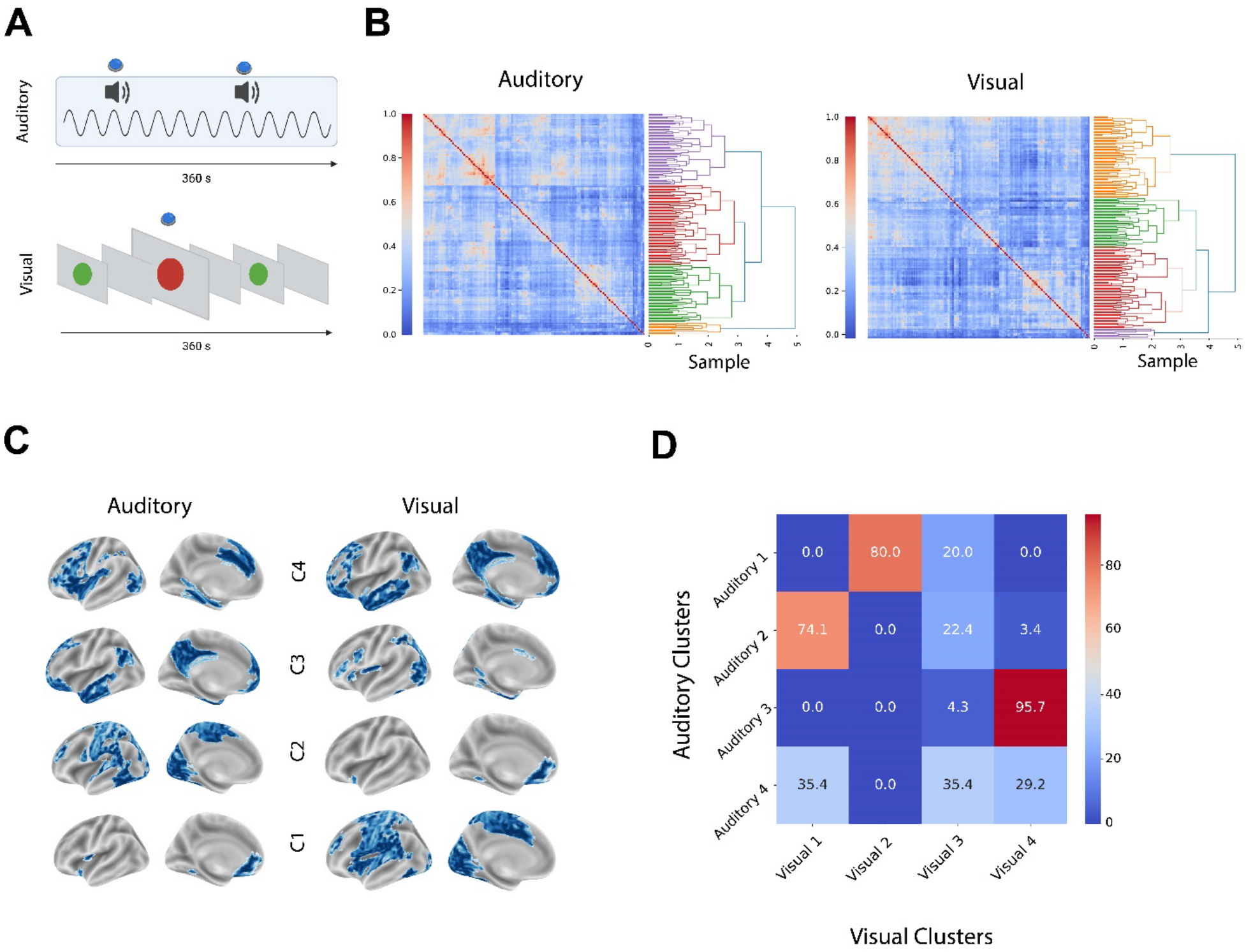
Functional connectivity profile of change detection. **A**, Overview of the experimental paradigm. Participants completed auditory and visual oddball detection tasks during simultaneous fMRI-EEG acquisition. Across three runs, total 375 stimuli (125 per run) were presented for 200 ms each, with a 2–3 s uniformly distributed ITI and a target probability of 0.2. In the auditory task, the standard was a 390 Hz pure tone and the target a broadband “laser-gun” sound. In the visual task, the standard was a small green circle and the target a large red circle, both on isoluminant gray backgrounds (1.15 and 3.45 DVA). **B**, Functional connectivity (FC) matrices (task’s left panel) were computed for auditory and visual runs by applying correlation coefficient to the fMRI time series across brain regions. Hierarchical clustering (task’s right) grouped regions with similar FC profiles, resulting in four clusters per modality. **C**, Brain maps show the spatial distribution of auditory and visual clusters, each reflecting functionally grouped regions based on FC patterns. **D**, Cluster overlap analysis revealed strong cross-modal similarity: auditory cluster 3 and visual cluster 4 shared 95% of regions; auditory cluster 1 and visual cluster 2 overlapped 80%; and auditory cluster 2 and visual cluster 1 overlapped 74%. ITI: Inter Trial Interval. DVA: Degrees of Visual Angle.

Visual inspection of mean FC matrices revealed organized and modular patterns of connectivity revealing groups of hierarchical networks and subnetworks with similar but also distinct features between the modalities. A strong diagonal structure was present for both modalities, reflecting higher connectivity among anatomically or functionally related regions. In the auditory condition, stronger connectivity was presented between temporal and posterior regions, consistent with engagement of auditory and multisensory systems. In contrast, the visual condition showed a more distinctly modular structure, with enhanced connectivity among occipital and posterior parietal regions involved in visual processing. The visual FC matrix appeared to exhibit greater functional segregation, suggesting more differentiated subnetworks engaged during visual change detection.

To further characterize these patterns, hierarchical clustering was applied to the group-level FC matrices. Four clusters of cortical regions with similar connectivity profiles were identified for each modality (Fig. 1C), and their spatial distributions were projected onto cortical surfaces for visualization.

Auditory cluster 1 and visual cluster 2 included regions in the inferior frontal gyrus and ventromedial prefrontal cortex, areas commonly associated with executive control, language processing, decision-making, and value representation^18,19^. These clusters likely reflect engagement of frontal control networks supporting integrative and regulatory functions during task performance.

Auditory cluster 2 spanned both primary sensory and association areas, including primary visual, auditory, and somatosensory cortices, as well as adjacent temporal, parietal, and frontal regions. This widespread cluster suggests a network supporting early sensory processing and cross-modal integration. Visual cluster 1 exhibited broad overlap with this auditory cluster, encompassing a similar set of regions but with relatively more focused engagement of dorsal parietal and medial occipital areas, and reduced extension into inferior temporal and anterior frontal regions.

Auditory cluster 3 and visual cluster 4 covered a set of regions including the posterior cingulate cortex, precuneus, retrosplenial cortex, inferior parietal lobule, and lateral prefrontal cortex. These areas are part of large-scale networks such as the default mode network (DMN), frontoparietal control network (FPN), and dorsal attention network (DAN)^20,21^. Their joint engagement suggests these clusters may support a combination of internally directed cognition, top-down attentional modulation, and cognitive control processes during task execution.

Auditory cluster 4 was localized primarily in occipitotemporal and ventral parietal cortices, including the fusiform gyrus, lateral occipital areas, inferior temporal cortex, lingual gyrus, and posterior parahippocampal cortex, regions implicated in high-level visual perception, scene recognition, and memory-guided processing^22–24^. Visual cluster 3 showed spatial overlap with this cluster but was more spatially restricted and discontinuous. It also included the posterior superior temporal sulcus and adjacent temporoparietal areas, indicating potential involvement in multimodal perceptual integration.

To assess cross-modal similarities, we quantified the proportion of spatial overlap between clusters across conditions by calculating the percentage of parceled areas that present in both clusters. The greatest overlap was observed between auditory cluster 3 and visual cluster 4, suggesting that this cluster reflects a modality-independent network shared across auditory and visual change detection. Additional overlaps were found between auditory cluster 1 and visual cluster 2 (80%), and between auditory cluster 2 and visual cluster 1 (74%). These findings indicate that while some connectivity patterns are modality-specific, others reflect common functional networks consistently engaged across sensory domains.

### 2.2. Latent connectivity structure reveals dual network organization and modality-sensitive patterns

Given the observed similarities in clustering-based FC patterns across modalities, we next investigated whether a shared latent structure underlies connectivity in auditory and visual change detection. To this end, we applied partial least squares (PLS) regression to decompose the covariance between the two mean auditory and visual FC matrices into a common low-dimensional subspace. A PLS model with four latent variables (LV) efficiently captured the shared variance structure, with component scores showing strong intermodal correlations and minimal variability between auditory and visual tasks (Fig. 2A), suggesting a robust cross-modal correspondence in FC architecture. To assess robustness to participant composition, we repeated the decomposition using leave-one-participant-out, independent split-half and participant-bootstrap resampling (Supplementary Fig. 1). Leave-one-out and bootstrap solutions closely reproduced the full-sample maps (median spatial r=0.998–0.999 and 0.963– 0.988, respectively). Across 1,000 non-overlapping split-half partitions, reproducibility remained high for LV1–LV2 (median r=0.930–0.949) and was more variable for LV3– LV4 (r=0.838–0.928), with the greatest variability for auditory LV4.

**Figure 2.**
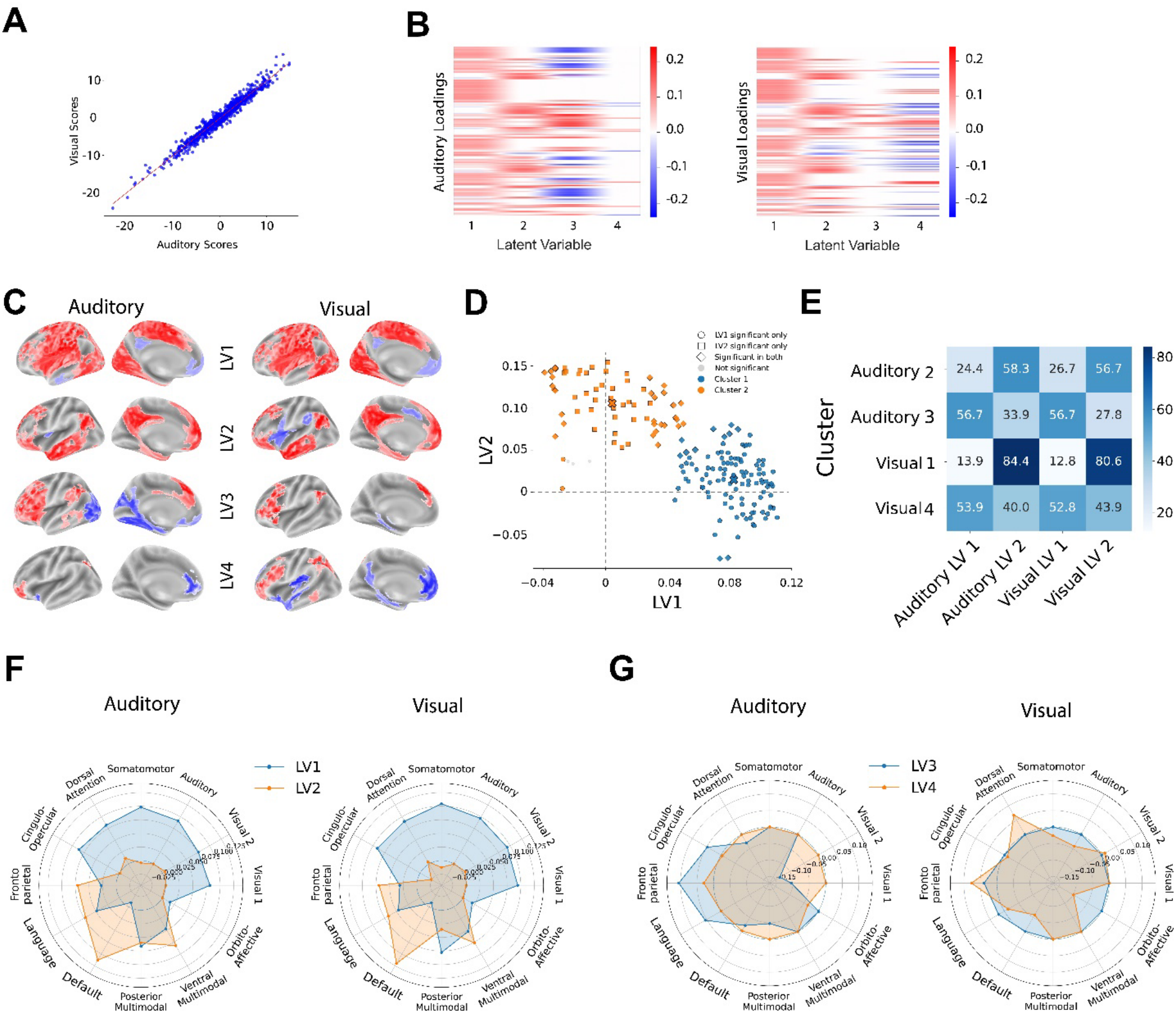
Latent connectivity structure reveals shared and modality-sensitive network modes. **A**, Partial least squares (PLS) regression was used to extract four latent variables (LVs) from the covariance between auditory and visual mean functional connectivity (FC) matrices. Component scores showed high cross-modal correlation, indicating shared latent structure. **B**, Regional loadings of LV1–LV4 in auditory and visual conditions. LV1 and LV2 showed non overlapping topographies across modalities, while LV3 and LV4 displayed modality-specific richness (auditory-dominant in LV3, visual-dominant in LV4) and bipolar patterns. Red indicates regions with positive loadings, contributing positively to the latent variable, while blue indicates negative loadings, contributing inversely. **C**, Surface projection of regions with significant contributions to each LV (p < 0.05, permutation test, 1,000 iterations), demonstrating distinct spatial patterns for each latent variable. Color scale reflects the same loading direction as in panel B. **D**, K-means clustering (k=2) applied to LV1 and LV2 loadings revealed two distinct regional groupings, supporting the presence of dual functional networks. **E**, Heatmap quantifying the percentage overlap between FC-derived clusters and regions with high LV loadings. Visual cluster 1 and auditory cluster 2 showed greatest overlap with LV2, indicating convergence on a modality-independent functional network. **F**, Projection of LV1 and LV2 loadings onto the 12-network CAB-NP atlas revealed conserved cross-modal organization, with LV1 emphasizing sensory–attention systems and LV2 showing greater default-mode, frontoparietal and multimodal involvement. **G**, Modality-biased LV3 and LV4 showed stronger frontoparietal contributions, with auditory LV3 and visual LV4 exhibiting distinct higher-order sensory–attentional profiles.

To test whether the reciprocal PLS solutions represented corresponding latent dimensions despite directional fitting, we compared them with a symmetric decomposition of the auditory–visual cross-covariance. Components matched one-to-one across formulations (LV1–LV4), with absolute spatial correlations of 0.970–1.000 for auditory and 0.976–1.000 for visual loadings; the corresponding four-dimensional subspaces were nearly identical (maximum principal angle <0.5°). The symmetric spectrum was strongly low-dimensional, with the first four components accounting for 97.1% of cumulative cross-modal structure (Supplementary Fig. 2).

To characterize the anatomical contributions to each LV, we computed regional loadings and assessed their significance using a nonparametric permutation test (1,000 iterations). Regions with significant contributions (p < 0.05, uncorrected) were visualized on the cortical surface (Fig. 2B–C). The first two components, LV1 and LV2, exhibited strikingly distinct spatial profiles with minimal anatomical overlap, implying they encode functionally separable network patterns. Participant-bootstrap resampling further supported the stability of these spatial patterns: 50–76% of parcel loadings had 95% bootstrap confidence intervals excluding zero, and 96.4–100% of permutation-selected parcels were recovered within the corresponding bootstrap-stable maps (Supplementary Fig. 3).

Furthermore, unsupervised K-means clustering (k = 2) closely recapitulated the permutation-defined separation of LV1- and LV2-selective regions (agreement = 0.99, adjusted Rand index = 0.94, permutation p = 2×10^-4^; Fig. 2D), demonstrating that the two LVs recruit highly separable sets of cortical regions in loading space. Notably, LV2 showed high spatial overlap with visual cluster 1 (80%) and auditory cluster 2 (84%) identified earlier via hierarchical clustering, suggesting that this latent component corresponds to a cross-modal, modality-independent network. Additional overlaps were observed between auditory cluster 2 and visual LV2 (56%), and between auditory cluster 3 and both LV1 components (53%), reinforcing the notion that the latent structure captured by PLS reflects functional motifs embedded within the original FC matrices (Fig. 2E).

To explore the functional architecture of these latent dimensions, we projected LV1 and LV2 onto the Cole-Anticevic Brain-wide Network Partition (CAB-NP) functional atlas^25^. This mapping revealed a conserved network-level organization across auditory and visual modalities. LV1 preferentially engaged sensory and externally oriented control systems, with strong contributions from visual, somatomotor, auditory, dorsal attention and cingulo-opercular networks. By contrast, LV2 was shifted toward higher-order associative systems, with prominent contributions from default, language, frontoparietal and multimodal networks and comparatively weaker representation of primary sensory networks (Fig. 2F). Notably, frontoparietal involvement was evident in both LVs, suggesting that their separation is not driven by a simple presence-versus-absence of executive control engagement, but rather by the broader network context in which frontoparietal regions participate. Together, these profiles distinguish an LV1 configuration emphasizing sensory, attentional and action-related processing from an LV2 configuration emphasizing associative, contextual and multimodal integration.

We next examined latent variables 3 and 4 to assess whether they captured modality-biased patterns. LV3 exhibited a richer and more differentiated topography in the auditory condition, while LV4 was more structured in the visual condition, suggesting each component captured a modality-dominant network mode (Fig. 2B). This asymmetry was preserved under participant-bootstrap thresholding, with greater spatial extent for auditory than visual LV3 and the opposite pattern for LV4 (Supplementary Fig. 4). Both components contained pronounced positive and negative loading patterns, indicating opposing contributions across distributed cortical systems. This configuration reflects functionally antagonistic relationships across the cortex, consistent with large-scale network opposition such as that observed between task-positive systems and the default mode network^26^.

In contrast to the modality-invariant dimensions, LV3 and LV4 exhibited distinct modality-dependent network profiles when projected onto the 12-network CAB-NP atlas (Fig. 2G). The auditory-biased LV3 showed its strongest positive expression within the frontoparietal network, accompanied by contributions from cingulo-opercular and language-related systems. By comparison, the visual-biased LV4 was characterized by prominent dorsal-attention and frontoparietal weighting, consistent with enhanced engagement of visuospatial selection and higher-order control processes. Thus, the modality-biased components were not dominated simply by their corresponding sensory networks; rather, both preferentially recruited frontoparietal control architecture within modality-specific network configurations. This organization suggests that sensory-context specificity during change detection may arise through differential configuration of higher-order control systems, with auditory processing preferentially coupled to frontoparietal–cingulo-opercular circuitry and visual processing to frontoparietal–dorsal-attention circuitry^27^.

### 2.3. Cross-modal connectivity profiles of latent variables

To characterise the inter-regional organisation of each latent variable (LV) and compare modalities, we constructed weighted graphs on a 180-region Glasser parcellation for every LV separately in the auditory and visual conditions. Nodes were retained when a region’s loading on the LV was significant. We applied an adaptive threshold (w>μ+kσ; k=1.5), which, under approximately normal weights, retains the strongest ≈7% of edges; findings were robust to k ∈ [1.0,2.0] and to density-matched proportional thresholds. Thus, each graph was node and edge-thresholded per LV and modality (Fig. 3A). We first compared regional connectivity profiles for LV1 and LV2 using degree centrality. Two-sample Kolmogorov–Smirnov (KS) tests revealed no differences between auditory and visual degree distributions for LV1 (KS = 0.028, P > 0.05) or LV2 (KS = 0.064, P > 0.05) (Fig. 3B). Edgewise overlap was high for LV1 (97%) and slightly reduced for LV2 (85%), indicating a conserved architecture with modest modality-specific tuning. For subsequent analyses, core networks of the graphs were defined as the top 15% highest-degree nodes within each graph.

**Fig. 3.**
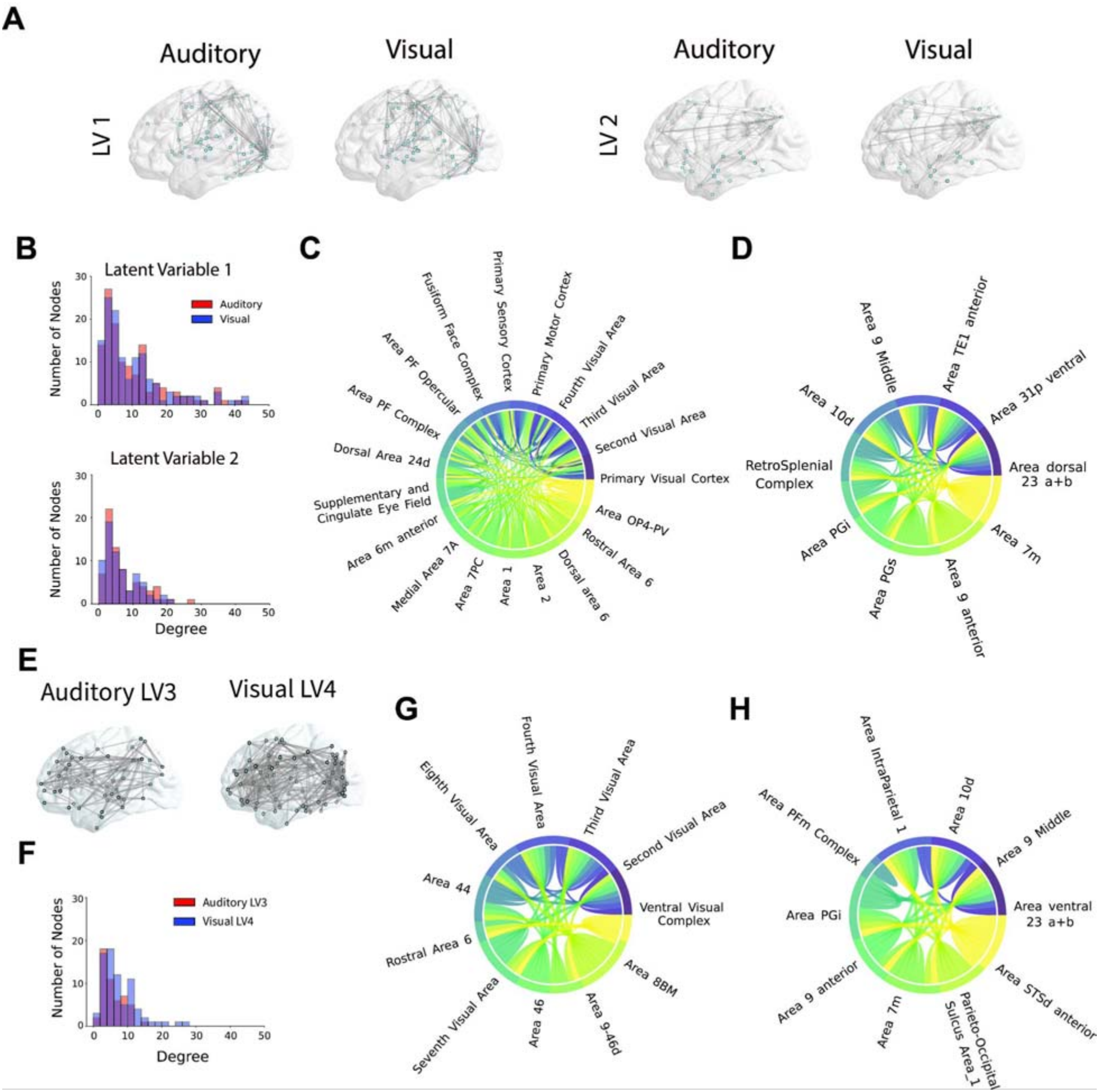
Cross-modal connectivity profiles of latent variables. **A**, Node- and edge-thresholded graphs (180-region parcellation) for LV1 and LV2 in the auditory and visual datasets. Edges were retained using an adaptive cutoff algorithm (see Methods). **B**, Degree distributions by modality; two-sample Kolmogorov–Smirnov tests showed no differences for LV1 (KS = 0.028, P>0.05) and LV2 (KS = 0.064, P>0.05). Edge overlap across modalities was 97% for LV1 and 85% for LV2. Core networks were defined as the top 15% highest-degree nodes. **C**, LV1 core network forms a contiguous sensory-to-action pathway: early visual cortex (Primary, Second, Third, Fourth Visual Areas) → posterior parietal/somatosensory nodes (Medial Area 7A, Area 7Pc, Areas 1–2, OP4–PV, PF opercular/complex, Primary Sensory Cortex) → premotor fields (rostral/dorsal Area 6, Area 6m anterior, Supplementary & Cingulate Eye Field) → Primary Motor Cortex. **D**, LV2 core network couples rostral/dorsolateral PFC (Area 10d, Area 9 Middle, Area 9 anterior), posterior cingulate/retrosplenial hubs (Area dorsal 23, Area 31p ventral, Retrosplenial Complex), medial/inferior parietal association cortex (Area 7m, Area PGs) and anterior temporal cortex (Area TE1 anterior); the only modality-specific node was Area PGi (visual) versus Area 8 lateral (auditory). **E**, Modality-biased LVs: auditory LV3 and visual LV4 graphs. **F**, Degree distributions were similar (KS = 0.166, P>0.05), but only 23% of edges overlapped, indicating distinct wiring. **G**, Auditory LV3 core network: widespread negative loadings in visual cortex (Second, Fourth, Seventh, Eighth Visual Areas; Ventral Visual Complex) alongside positive loadings in a frontal–premotor set (Area 46, Area 9-46d, Area 8BM, Area 44, rostral Area 6), consistent with auditory-centered control and visual suppression. **H**, Visual LV4 core network: predominantly loadings with negative contribution across rostral PFC, posterior cingulate/precuneus, STS and IPL (Area PGi), with positive loadings confined to intraparietal (Area IntraParietal 1) and PFm nodes, consistent with visual attentional sharpening and response preparation.

In LV1, the core network formed a contiguous sensory-to-action pathway: early visual cortices (Primary Visual Cortex; Second, Third and Fourth Visual Areas) feed posterior parietal nodes (Medial Area 7A, Area 7Pc, Areas 1–2) and opercular/primary somatosensory regions (OP4–PV, PF opercular/complex, Primary Sensory Cortex), which converge on premotor fields (rostral and dorsal Area 6, Area 6m anterior, Supplementary & Cingulate Eye Field) and Primary Motor Cortex (Fig. 3C). Functionally, this configuration implements a low-latency “detect-and-respond” pipeline: early visual areas register feature change^28^; parietal cortex encodes salience and spatial mapping^29^; SCEF supports orienting^30^; premotor regions set motor readiness^31^; and motor/somatosensory cortices execute the button press with rapid feedback^32^. The inclusion of fusiform cortex is consistent with ventral perceptual processing when visual input is present^33^, yet the backbone is not visual-specific; the near-identical architecture across modalities indicates a supramodal sensorimotor scaffold that routes both auditory- and visual-change signals into the same premotor–motor channel. Observing this same LV1 configuration in both visual and auditory modalities suggest that LV1 captures a supramodal “detect-and-respond” circuit. In short, LV1 reflects a modality-invariant, low-latency sensorimotor pipeline that links stimulus changes to orienting, motor preparation, and execution, precisely the operations required by this task.

LV2 core network captured rostral/dorsolateral prefrontal cortex (Area 10d, Area 9 middle, Area 9 anterior), anterior temporal association (Area TE1 anterior), posterior cingulate/retrosplenial hubs (Area dorsal 23, Area 31p ventral, Retro Splenial Complex), and medial/inferior parietal association cortex (Area 7m, Area PGs) (Fig. 3D). This constellation is characteristic of high-level control and context integration: rostral/dorsolateral PFC maintains and updates task sets^34^; posterior cingulate/retrosplenial regions provide internal context and confidence/evidence evaluation^35^; parietal association areas integrate multimodal inputs with current policies^36^; TE1 anterior contributes associative content^37^. Modality-specific tuning was minimal: the visual network additionally recruited Area PGi (inferior IPL), consistent with posterior IPL–based evidence integration, whereas the auditory network included Area 8 lateral (lateral BA8), consistent with enhanced top-down attentional selection and response gating. In contrast to LV1, LV2 reflects a modality-general control layer that maintains task context, integrates evidence and regulates attentional and response states^38^.

Because LV3 and LV4 exhibited modality-biased expressions (auditory-rich and visual-rich, respectively), we focused on auditory LV3 and visual LV4 (Fig. 3E). Degree distributions were similar (KS = 0.166, P > 0.05; Fig. 3F), yet only 23% of edges overlapped, indicating comparable hubness but distinct wiring. For LV3 (auditory), the core network showed widespread negative weights across visual cortex (Second Visual Area, Third Visual Area, Fourth Visual Area, Seventh Visual Area, Eighth Visual Area and Ventral Visual Complex) coupled with positive weights in a compact frontal– premotor set (Area 46, Area 9-46d, Area 8BM, Area 44, Rostral Area 6) (Fig. 3G). The signed structure indicates a control state that suppresses visual representations while up-regulating dorsolateral/medial frontal control and premotor readiness, consistent with cross-modal attentional focusing on the auditory stream: lateral/medial prefrontal nodes (Area 46, Area 9-46d, Area 8BM) sustain task set and bias sensory gain^34^; Area 44 supports selection/inhibition of competing inputs^39^; Rostral Area 6 prepares the motor response^40^. Negative occipitotemporal contributions suggest reduced visual interference, while LV3 reflects an auditory-biased control state supporting task maintenance, attentional selection and response preparation.

For LV4 (visual), loadings were predominantly negative across associative prefrontal and medial/posterior hubs (Area 10d, Area 9 Middle, Area 9 anterior, Area ventral 23, Area 7m L, Parieto-Occipital Sulcus Area 1, Area STSd anterior, Area PGi), with positive weights restricted to Area IntraParietal 1 and Area PFm Complex (Fig. 3H). This pattern indicates that greater LV4 expression corresponds to relative down-weighting of rostral PFC, retrosplenial/posterior cingulate and high-level temporal/Inferior parietal representations, while amplifying a dorsal attention/sensorimotor selection axis centered on intraparietal and PFm nodes. Functionally, LV4 is consistent with a visual attentional sharpening state: dorsal intraparietal cortex prioritizes spatial selection and visuomotor mapping^29^, PFm supports stimulus–response coupling^41^, and broad associative/contextual systems are concurrently suppressed to reduce interference^42^. In the visual change-detection task, LV4 therefore reflects a mode that biases processing toward rapid target selection and response readiness in visual space, achieved by strengthening dorsal parietal control while attenuating elaborative prefrontal, retrosplenial and STS processing.

### 2.4 Temporal latent variables reveal shared latent dynamics and modality-dependent target trajectories

To examine the temporal expression of the connectivity dimensions identified by PLS, we projected parcel-wise BOLD activity onto the corresponding PLS loading vectors to derive four temporal latent variables (tLVs) for each run (Fig. 4A; Methods). tLV1 and tLV2 showed the largest and most slowly varying fluctuations, whereas tLV3 and tLV4 showed smaller-amplitude dynamics (Fig. 4B). Spectral analysis confirmed greater low-frequency power for tLV1 and tLV2 than for the remaining components (Supplementary Fig. 5). The temporal geometry paralleled the spatial PLS organization. Auditory and visual observations overlapped strongly in both latent projections but showed different covariance orientations in the tLV3–tLV4 plane, with auditory variability extending preferentially along tLV3 and visual variability along tLV4 (Fig. 4C). By contrast, covariance structure in the tLV1–tLV2 plane was closely aligned across modalities. Thus, the dominant temporal dynamics were expressed in a shared tLV1–tLV2 space, with modality-related variation more apparent along the secondary dimensions.

**Fig. 4.**
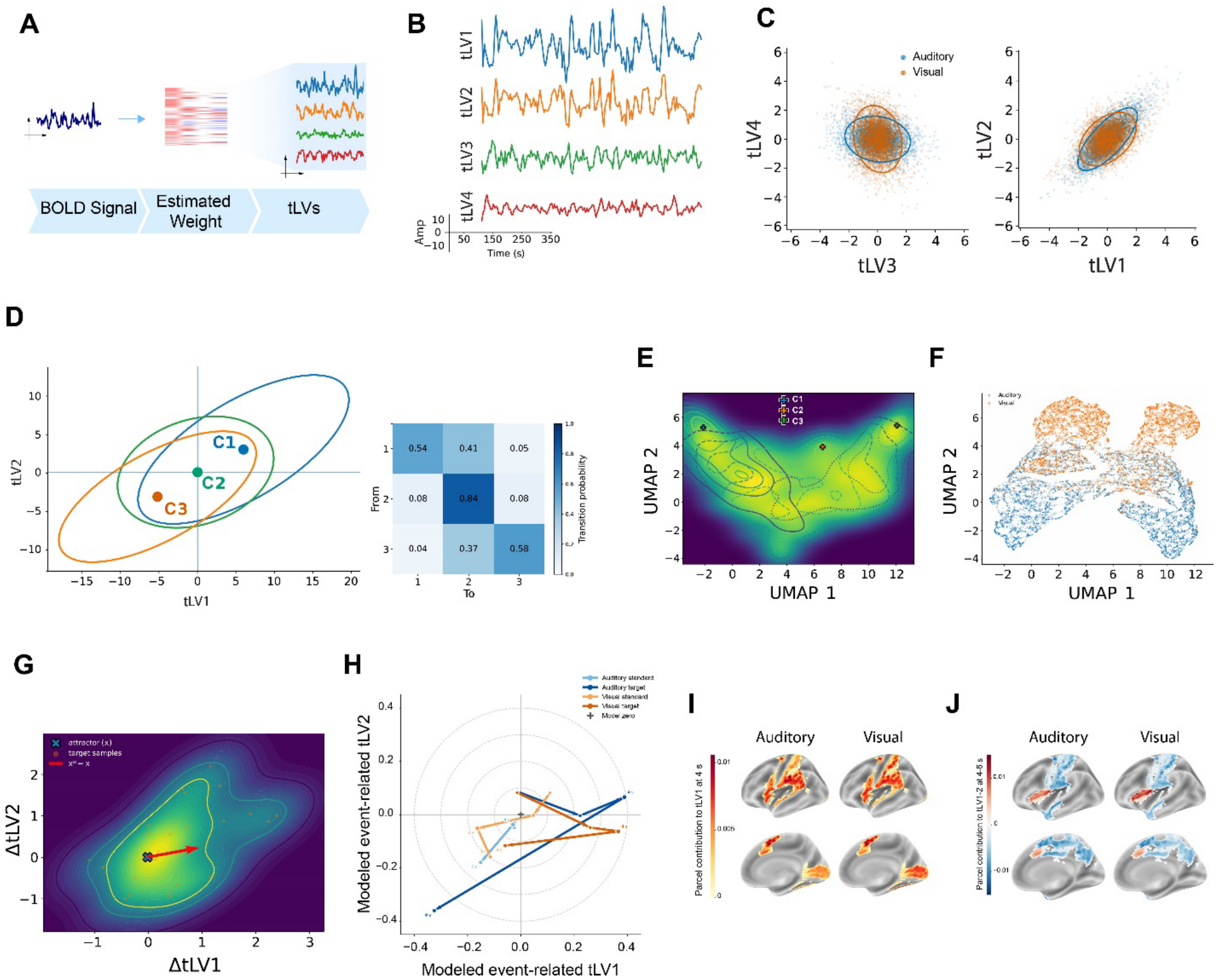
Shared latent organization and modality-dependent target-associated dynamics. **A**, Schematic of temporal latent-variable derivation. Parcel-wise BOLD time series were weighted by their corresponding PLS loadings and summed across parcels to generate a tLV time course for each component. **B**, Representative run illustrating the time-resolved expression of four tLVs. tLV1 and tLV2 exhibited larger-amplitude, slowly varying fluctuations, whereas tLV3 and tLV4 showed smaller-amplitude fluctuations. **C**, Modality-related organization of latent space. Auditory and visual observations overlapped in both projections. In the tLV3–tLV4 plane (left), covariance ellipses differed in orientation, with auditory variability extending more strongly along tLV3 and visual variability along tLV4. In the tLV1–tLV2 plane (right), covariance structure was more closely aligned across modalities. **D**, Three Gaussian mixture model (GMM)-defined latent configurations and their transition probabilities. A three-component GMM was fitted to the pooled four-dimensional tLV observations, with component centroids and covariance ellipses shown in the tLV1–tLV2 projection (left). Components were designated positive (C1), central (C2) and negative (C3) configurations. The row-normalized transition matrix (right) shows the highest persistence in the central configuration (P = 0.84), with infrequent direct transitions between the peripheral configurations. **E**, Two-dimensional UMAP embedding of the pooled four-dimensional tLV observations. The background shows pooled observation density, while GMM posterior-weighted density contours and their peaks indicate smoothly connected regions of preferential configuration occupancy. **F**, UMAP embedding of tLV3–tLV5 derived from a separate five-component PLS decomposition, excluding the two dominant shared dimensions. Auditory and visual observations showed partially differentiated distributions while retaining substantial overlap, revealing modality-related structure within the higher-order latent dimensions. **G**, Model-based characterization of target-associated displacement in tLV1–tLV2 space. The density field shows the empirical distribution of target-associated observations relative to the baseline fixed-point estimate of the second-order vector autoregressive model (cross); the arrow shows the model-predicted displacement under target input, oriented predominantly along positive tLV1. **H**, Overlap-adjusted event-related trajectories estimated using a finite impulse response model in which target and standard events were modeled jointly. Auditory and visual target and standard responses are shown in the tLV1–tLV2 plane across post-event lags. At 4 s, targets were displaced positively from standards along tLV1 in both modalities, followed by divergent 4–6-s trajectories, most prominently along tLV2. **I**, Parcel-wise contributions to the target–standard tLV1 contrast at 4 s. The strongest positive contributors were selected separately in auditory and visual runs and displayed using their union and a common color scale, revealing a largely overlapping anatomical contribution pattern across modalities. **J**, Parcel-wise contributions to the 4–6-s evolution of the target–standard contrast. Twenty-seven parcels were selected jointly from the strongest tLV1 and tLV2 contributors, with each parcel displaying its assigned latent-dimension contribution using the same assignment across modalities. Auditory runs showed a stronger negative contribution pattern than visual runs.

We next characterized temporal dependence within the dominant tLV1–tLV2 subspace using vector autoregressive models. A second-order model provided a compact description of the dynamics (cross-validated R2=0.43±0.16). Including target timing as an external input produced only a small improvement in predictive performance (R2=0.440.16; ΔR2=0.005±0.015), indicating that most temporal variance was captured by the ongoing latent dynamics. The resulting dynamical landscape revealed a smooth, unified manifold in tLV1–tLV2 space, organized around a central stability (Supplementary Fig. 6). Extending the model to tLV3 and tLV4 did not materially improve prediction or alter the dominant geometry.

To describe preferential occupancy within this continuous latent space, we fitted a three-component Gaussian mixture model to the pooled four-dimensional tLV trajectories (17,170 observations; Fig. 4D). Components were ordered by their projected tLV1–tLV2 centroid coordinates and termed positive, central and negative latent configurations. The central configuration accounted for 70.4% of observations, whereas the positive and negative configurations accounted for 14.7% and 15.0%, respectively. Assignments were frequently probabilistic rather than categorical: 57.7% of observations had a maximum posterior probability below 0.8. Accordingly, these configurations are best interpreted as regions of preferential occupancy within a continuous latent distribution rather than discrete neural states.

Transition probabilities were consistent with this organization (Fig. 4D). The central configuration showed the greatest persistence (C2 P=0.84), whereas the positive and negative configurations were less persistent (C1 P=0.54 and C3 P=0.58) and transitioned preferentially toward the central configuration (C1→2 P=0.41 and C3→2 P=0.37, respectively). Direct transitions between the two peripheral configurations were uncommon (C1→3 P=0.05 and C3→1 P=0.04). Projection of the full four-dimensional trajectories into UMAP space likewise revealed overlapping, smoothly connected configuration-weighted density fields rather than isolated clusters (Fig. 4E). Furthermore, auditory and visual time points occupied a largely overlapping manifold (Supplementary Fig. 7).

To examine modality-related organization independently of the two dominant shared dimensions, we repeated the PLS decomposition with five components and projected tLV3–tLV5 into a separate UMAP embedding (Fig. 4F). In this representation, auditory and visual observations showed more differentiated distributions, with visual observations concentrated toward the upper portions of the embedding and auditory observations toward the lower portions, while retaining substantial overlap in the central regions. Thus, although the dominant latent architecture was largely shared across modalities, the higher-order dimensions revealed additional modality-related structure within the temporal expression of cortical activity.

We then asked how target events were expressed within this shared latent organization. In the VAR model, target input produced a predicted displacement predominantly along positive tLV1, with comparatively little displacement along tLV2 (Fig. 4G). We therefore estimated empirical target- and standard-associated trajectories using a joint finite impulse response model in which lagged regressors for target and standard events were fitted simultaneously. Because stimuli occurred every 2–3 s, this formulation accounted for temporal overlap between the hemodynamic responses to adjacent events rather than treating each event in isolation. Target–standard separation was greatest along tLV1 at 4 s in both modalities (Fig. 4H), reaching +0.418 within-run SD in auditory runs and +0.528 SD in visual runs (Holm-adjusted P≈3.1×10-5 in both; exploratory). Thus, auditory and visual targets initially displaced activity along a closely shared latent direction.

The regional contributions to this 4-s tLV1 displacement were also strongly overlapping across modalities (Fig. 4I). Prominent positive contributors included the supplementary and cingulate eye field, dorsal and rostral premotor cortex, primary somatosensory cortex, and parietal and frontal opercular cortices. Modality-related weighting was superimposed on this common pattern: auditory runs showed relatively stronger contributions from auditory and superior temporal regions, including the Auditory 4 complex and temporal area 2, whereas visual runs showed stronger contributions from visual cortex and the ventral visual complex. These maps therefore identify a shared set of premotor, somatosensory and opercular contributors to the early target-associated tLV1 response, together with sensory-weighting differences expected from the two task modalities.

The trajectories diverged more clearly between 4 and 6 s (Fig. 4H). This divergence was most apparent along tLV2: auditory trajectories shifted in the negative direction (−0.283 SD), whereas visual trajectories showed little net change and remained slightly positive (+0.053 SD). Parcel decomposition indicated that the auditory shift reflected a different balance of opposing regional contributions rather than recruitment of an entirely distinct anatomical set. The strongest negative tLV2 contributors showed a larger combined negative contribution in auditory than visual runs (exploratory exact sign-flip P=0.0086, Holm-adjusted P=0.0345), whereas the corresponding positive-contributor comparison was not significant.

To summarize the anatomical organization of this late divergence across both dominant dimensions, we jointly selected the strongest contributors from tLV1 and tLV2, yielding 27 unique parcels (Fig. 4J). These included premotor and somatosensory cortex, supplementary and cingulate eye field, opercular and peri-Sylvian regions, medial parietal and cingulate cortex, and superior temporal association areas. The same parcel-to-LV assignment was retained across auditory and visual maps for visualization. Although the anatomical distribution was broadly shared, the signed contribution pattern differed between modalities, with auditory runs showing a stronger negative component, particularly along tLV2, and visual runs showing a more balanced contribution pattern.

Consistent with this observation, the jointly selected parcels differed between modalities in their two-dimensional tLV1–tLV2 contribution vectors (exploratory exact sign-flip P=4.6×10-5, Holm-adjusted P=1.37×10-4), whereas their overall vector magnitudes did not differ reliably. Thus, the late auditory–visual divergence was expressed primarily as a redistribution of contributions across shared latent dimensions rather than a difference in total response strength. Together, these analyses identify a common early target-associated displacement along tLV1 followed by modality-dependent evolution of the same low-dimensional cortical architecture.

## 3 Discussion

Here we show that auditory and visual change detection are organized within a shared latent connectivity and dynamical space. Auditory and visual target detection involved closely related patterns of large-scale connectivity, yet their temporal responses were not identical. Across modalities, two dominant latent dimensions captured a shared organization of sensory, attentional, motor and associative systems, whereas additional dimensions retained modality-biased network structure. This correspondence extended to ongoing BOLD dynamics, with auditory and visual activity occupying overlapping regions of the same low-dimensional space. Targets were initially associated with a similar displacement along tLV1 in both modalities, but the trajectories subsequently diverged, particularly along tLV2. These findings suggest that common network organization need not imply identical temporal processing. Instead, auditory and visual tasks appear to draw on a largely shared cortical architecture while differing in how its constituent regions contribute over the course of a target-associated response.

Previous oddball and change-detection studies have consistently identified responses extending beyond primary sensory cortex, particularly in temporoparietal, frontal, insular, cingulate and premotor regions ^1–3^. The overlap of these responses across modalities has supported the idea of a supramodal system for detecting behaviorally relevant events and preparing appropriate actions ^4,5,43^. At the same time, more recent work has shown that modality-specific and supramodal responses coexist, with their relative contributions depending on stimulus properties, task demands and the temporal structure of sensory expectations ^8–10^. Regional overlap alone, however, cannot establish whether two tasks engage the same distributed network organization. The same cortical regions may participate in different patterns of coordination, just as distinct sensory pathways may converge on common attentional and motor systems. By examining connectivity profiles, latent loading patterns and their temporal expression, the present study provides a complementary account of how shared and modality-dependent contributions are organized within the same functional architecture.

The two dominant PLS dimensions captured different aspects of this shared organization. LV1 was weighted toward visual, auditory, somatomotor, dorsal-attention and cingulo-opercular systems, with additional frontoparietal contributions. Its core network included sensory and posterior parietal regions alongside somatosensory, opercular and premotor cortex. This pattern is consistent with the coordination of sensory information, attentional selection and response preparation during active target detection ^4,29,32,43^. LV2, in contrast, showed greater contributions from default-mode, language, frontoparietal and multimodal association networks, including prefrontal, posterior cingulate, retrosplenial and parietal association regions. Prefrontal regions are implicated in maintaining and updating task goals ^34,38^, whereas posterior cingulate and default-mode systems have been associated with contextual and internally oriented processing ^35,44^. These contributions suggest that LV2 may support the integration of sensory information with ongoing task context. Importantly, frontoparietal regions contributed to both dimensions, indicating that their distinction lies less in the involvement of separate control systems than in the broader networks with which these regions are coordinated. Whereas LV1 emphasizes sensory, attentional and motor processing, LV2 is more closely associated with contextual and integrative functions. Together, these dimensions suggest that the widespread cortical responses observed during target detection may reflect complementary aspects of processing organized within a shared network architecture.

The two remaining latent variables revealed how sensory modality was represented within this common organization. Auditory LV3 combined positive loadings in lateral and medial prefrontal and premotor regions with negative loadings across occipitotemporal visual areas. Its frontoparietal and cingulo-opercular weighting is consistent with the involvement of frontal systems in attentional selection and cognitive control ^39^. Visual LV4 showed a different pattern, with positive contributions concentrated in intraparietal and inferior parietal regions, which have established roles in visuospatial selection ^41^, and negative contributions distributed across prefrontal, posterior medial and temporal association cortex. These differences are consistent with evidence that frontoparietal regions can carry modality-dependent information despite their broader involvement in domain-general control ^27^. Although the opposing loading signs do not establish active suppression between networks, they reveal distinct patterns of contribution across sensory and control-related regions. Modality bias was therefore evident not only in sensory cortex but also in the organization of distributed association and control systems.

The graph analyses provided a complementary view of these differences. The dominant dimensions showed highly conserved auditory–visual connectivity organization, with 97% edge overlap for LV1 and 85% for LV2, consistent with their shared spatial loading profiles. Auditory LV3 and visual LV4, by contrast, had similar degree distributions but only 23% edge overlap. Comparable numbers of connections per region therefore concealed substantial differences in which regions were connected. This distinction matters because a similar distribution of regional connectivity does not necessarily imply the same network architecture ^45–48^. The graph results strengthen the interpretation that LV1 and LV2 capture conserved patterns of inter-regional coordination, while the modality-biased dimensions are characterized by different arrangements of connections among sensory, associative and control-related regions.

Projecting the PLS dimensions onto time-resolved BOLD activity revealed a close correspondence between the spatial organization of connectivity and its temporal expression. Previous work has shown that task-dependent fluctuations in functional connectivity occur within a relatively stable large-scale architecture ^49,50^, while recurrent patterns of network integration accompany ongoing cognitive performance ^51^. In the present data, tLV1 and tLV2 captured the largest low-frequency fluctuations and showed closely aligned covariance geometry across auditory and visual runs. By contrast, modality-related differences were more apparent along tLV3 and tLV4, extending the distinction between shared and modality-biased connectivity dimensions into the temporal domain. The distribution of observations within latent space was also structured, with three partially overlapping Gaussian-mixture configurations and a predominance of central configuration occupancy. Unlike discrete-state interpretations of cognitive dynamics ^15^, these configurations did not exhibit sharp boundaries but described graded regions of preferential occupancy within a continuous distribution. This organization suggests that auditory and visual activity follows similar dominant latent trajectories while preserving modality-related variation in their secondary dimensions.

The target-related analyses showed how behaviorally relevant events were expressed within this ongoing organization. Classical oddball studies have identified transient target-associated responses across frontal, parietal and cingulo-opercular systems, while simultaneous EEG–fMRI investigations have linked aspects of target evaluation to distributed cortical activity ^52,53^. Consistent with this literature, target input in our autoregressive model produced a predicted displacement predominantly along tLV1. The overlap-adjusted FIR analysis provided complementary empirical evidence, revealing a pronounced target–standard separation along the same dimension at 4 s in both modalities. This displacement was associated with positive contributions from the supplementary and cingulate eye field, premotor and somatosensory regions, and parietal and frontal opercular cortices. These areas overlap with systems implicated in attentional orienting and the preparation of goal-directed responses ^29–32^. Although auditory and visual targets differed in their sensory cortical contributions, both were associated with a common premotor–sensorimotor–opercular pattern. This finding is consistent with evidence for modality-general contributions to speeded sensory detection ^43^, while suggesting that the shared response is embedded within a broader latent architecture rather than confined to a fixed set of target-responsive regions.

The common displacement at 4 s was followed by a more pronounced auditory–visual divergence between 4 and 6 s, particularly along tLV2. Auditory trajectories shifted toward negative tLV2 values, whereas visual trajectories showed relatively little net change along this dimension. This difference was not explained by an entirely separate set of cortical contributors. Instead, the parcel-level analysis identified a different balance of positive and negative contributions across largely overlapping premotor, somatosensory, opercular, medial parietal and temporal association regions. The stronger negative tLV2 contribution in auditory runs was especially notable, while the jointly selected tLV1–tLV2 parcel vectors differed in their configuration without a reliable difference in overall magnitude. This distinction is relevant to previous accounts separating transient target-related processing from more sustained attentional control ^5^, as well as evidence that higher-order control regions can carry modality-dependent representations ^27^. Our findings suggest that the temporal evolution of target-associated activity may similarly differ between sensory modalities, even when the underlying connectivity architecture is largely conserved. Although the late interval was selected exploratorily and requires independent confirmation, the results indicate that cross-modal differences can emerge through changes in the relative contributions of shared cortical regions rather than through the recruitment of separate networks.

The interpretation of these trajectories is shaped by the demands of the oddball paradigm. Targets differed from standards in probability, physical properties, salience and response requirements, making it difficult to attribute the observed effects exclusively to sensory change detection. Although the joint FIR model accounted for overlapping hemodynamic responses, it could not disentangle target processing from the associated motor response. Paradigms that independently manipulate sensory deviance, behavioral relevance and response demands could help clarify these contributions. Moreover, the 4-s separation and subsequent 4–6-s divergence reflect modeled BOLD responses rather than the precise timing of neural events. The simultaneously acquired EEG could help relate these slower network dynamics to electrophysiological responses associated with target processing.

The network findings also require consideration of what the methods can establish. Functional connectivity and PLS describe statistical relationships rather than causal interactions ^54^, while opposing latent loadings do not necessarily indicate neural excitation or inhibition. The symmetric bilateral cortical parcellation further obscured hemispheric differences and excluded subcortical structures that may contribute to sensory selection and action preparation. Extending the analysis to the thalamus, basal ganglia and brainstem, particularly given previous evidence for brainstem involvement in oddball processing, could provide a more complete account of this network organization. Examining passive mismatch paradigms, other sensory modalities and tasks without motor responses would also help determine how broadly the shared latent architecture generalizes.

Together, these findings highlight the value of examining brain activity beyond individual regions and their responses, by considering how distributed cortical systems are organized and evolve over time. By combining dimensionality reduction of high-dimensional functional connectivity patterns with temporal analyses, we identified a largely conserved architecture across auditory and visual tasks, within which target-associated activity followed initially similar but subsequently modality-dependent trajectories. This approach links the spatial organization of large-scale networks to their ongoing and event-related dynamics, revealing differences in how shared cortical systems are expressed across sensory contexts. More broadly, it provides a framework for studying how complex brain activity can be understood through a smaller set of interpretable, dynamically evolving dimensions.

## Methods

### Dataset and participants

The present study reanalysed the publicly available Auditory and Visual Oddball EEG-fMRI dataset, originally reported by Walz et al. (2013)^55,56^. The dataset comprises simultaneous EEG and fMRI recordings acquired while healthy adults performed paradigm-matched auditory and visual oddball target-detection tasks. Only the fMRI data were analysed in the present study. The original sample included 17 healthy participants (6 female; mean age, 27.7 years). Each participant completed three auditory and three visual runs. The original study was approved by the Columbia University Institutional Review Board and all participants provided informed consent. One auditory run was excluded because of a technical acquisition problem, yielding 50 auditory and 51 visual runs (101 runs total) for analysis.

### Experimental paradigm

Participants performed separate but closely paradigm-matched auditory and visual oddball tasks. Each modality comprised three runs, with 125 stimuli per run and 375 stimuli across the three runs. Stimuli were presented for 200 ms, with an inter-stimulus interval uniformly distributed between 2 and 3 s. Targets occurred with probability 0.2 and standards with probability 0.8. Participants responded to target events with the right index finger using an MR-compatible response device.

In the auditory condition, the standard stimulus was a 390-Hz pure tone, and the target was a broadband “laser-gun” sound. In the visual condition, standards were small green circles presented on an isoluminant grey background and targets were larger red circles. The two tasks therefore shared the same target-detection structure while differing in sensory modality.

### MRI acquisition

MRI data were acquired on a 3-T Philips Achieva scanner. Whole-brain functional data were collected using an echo-planar imaging sequence with repetition time (TR) = 2,000 ms (i.e., sampling rate = 0.5 Hz), echo time (TE) = 25 ms, 32 slices, a 64 × 64 acquisition matrix, no inter-slice gap, and 3 × 3 × 4 mm voxels. Each functional run contained 170 volumes, corresponding to 340 s of acquisition. The original dataset also included higher-resolution images for registration and anatomical reference, as described in the source publication and OpenNeuro dataset documentation. Although EEG was recorded simultaneously with fMRI in the original experiment, the present analyses were restricted to the BOLD data.

### fMRI preprocessing

Raw BOLD data were preprocessed using fMRIPrep^57^; the exact minor software version was not retained. Data were submitted to the standard fMRIPrep functional preprocessing workflow, including estimation and correction of head motion, functional-to-anatomical alignment, and spatial normalization to MNI standard space. The MNI-normalized BOLD images underwent three additional image-level operations: temporal high-pass filtering at 0.01 Hz, brain extraction/masking, and spatial smoothing with a 2-mm Gaussian kernel. No additional nuisance-regression model was applied during the subsequent parcel-level time-series extraction.

### Cortical parcellation and regional BOLD time series

Cortical activity was represented using volumetric MNI-space implementation of the Human Connectome Project multimodal parcellation^17^. The original HCP-MMP1.0 atlas delineates 180 cortical areas per hemisphere. The atlas used here was a bilaterally symmetric 180-label volumetric representation, such that homologous left- and right-hemisphere territories shared the same parcel identifier. Each functional run was therefore represented by 180 regional time series. The mean signal within each parcel was standardized across time within each run using sample z-scoring. For parcel *i* in run *r*,

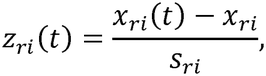

where *x*_*ri*_(*t*)is the parcel-averaged BOLD signal at time *t, x*_*ri*_ is its run-wise temporal mean, and *s*_*ri*_ is its sample standard deviation. This yielded a *T*_*r*_ × 180 standardized parcel-by-time representation for each run.

### Functional-connectivity estimation

Functional connectivity (FC) was estimated independently for each run as the Pearson correlation coefficient (range −1 to 1) between every pair of parcel time series. For parcels *i* and *j*,

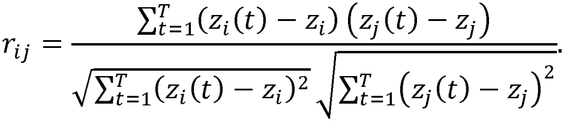

This produced a symmetric 180 × 180 FC matrix 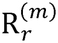 for run *r* and modality *m* ∈ {*A,V*}, where *A* and *V* denote auditory and visual conditions, respectively. Group-level auditory and visual FC matrices were obtained by averaging run-level matrices within each modality,

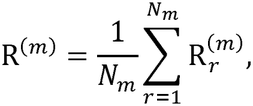

with *N*_*A*_ =50 and *N*_*V*_ =51. Correlation coefficients were averaged directly, matching the final implemented analysis; Fisher *r*-to-*z* transformation was not applied before group averaging (preliminary analysis demonstrated that the transformation did not significantly affect results in subsequent steps).

### Hierarchical organization of auditory and visual connectivity

To characterize coarse connectivity organization independently in the two sensory modalities, agglomerative hierarchical clustering was applied to the rows of each group-average FC matrix. Each cortical parcel was represented by its complete 180-dimensional connectivity profile,

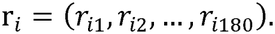

The distance between parcels was defined as Euclidean distance between their connectivity profiles,

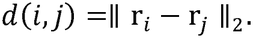

Clusters were merged using complete linkage, for which the distance between clusters *C*_*a*_ and *C*_*b*_ is

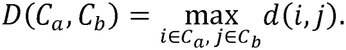

Optimal leaf ordering was used for dendrogram visualization. During exploratory analysis, the auditory and visual dendrograms were cut at linkage distances of 3.0 and 3.5, respectively, yielding four clusters in each modality. Cluster labels were subsequently renumbered according to dendrogram display order.

The four-cluster resolution was retained because it produced an interpretable coarse-grained organization in both modalities and enabled a common descriptive resolution for comparison with the four-dimensional PLS representation described below. Accordingly, the cluster number is treated as an analysis resolution rather than as evidence for a uniquely optimal biological partition.

### Cross-modal overlap of hierarchical clusters

Spatial correspondence between auditory cluster *A*_*i*_ and visual cluster *V*_*j*_ was quantified as the percentage of parcels in auditory cluster *i* that were also assigned to visual cluster *j*,

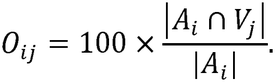

This measure is directional and row-normalized relative to auditory-cluster size. It is therefore not a Jaccard coefficient and need not be symmetric under exchange of auditory and visual clusters.

### Cross-modal partial least-squares decomposition

Partial least-squares (PLS) regression was used to identify low-dimensional structure linking auditory and visual FC organization. The two group-average FC matrices,

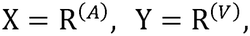

both had dimensions 180 × 180. In this formulation, the 180 observations corresponded to cortical parcels, and the 180 variables represented each parcel’s whole-brain connectivity profile. Thus, the decomposition operated on parcel-level group connectivity architecture, not on participant-level observations.

PLS derives latent score vectors

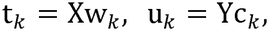

with weight vectors w_*k*_ and c_*k*_ chosen to maximize covariance between the corresponding projected representations. Conceptually,

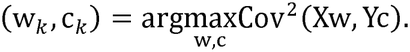

The matrices can be expressed as

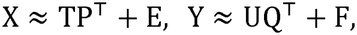

where T and U contain latent scores, P and Q contain loadings, and E and F are residual terms.

Four latent variables (LV1-LV4) were retained because this resolution provided a compact and mutually interpretable representation across the subsequent spatial, graph and temporal analyses, with dominant dimensions capturing cross-modal structure and later dimensions preserving modality-biased structure.

For the parcel-level score correspondence, auditory FC was entered as X and visual FC as Y, yielding auditory and visual score matrices T and U. Correspondence for component B was summarized by Pearson correlation,

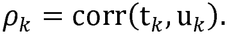

To obtain modality-specific response loading matrices while preserving a reciprocal cross-modal formulation, PLS was additionally fit in both directions. Auditory response loadings were obtained by predicting auditory FC from visual FC and retaining the response matrix. Visual response loadings were obtained by predicting visual FC from auditory FC and retaining the corresponding response-loading matrix.

### Symmetric cross-modal decomposition and component correspondence

To assess correspondence between reciprocal PLS solutions, the auditory–visual cross-covariance of standardized connectivity profiles was also decomposed symmetrically using singular-value decomposition (SVD). The paired singular vectors defined reference components against which the auditory and visual reciprocal response loadings were compared. Components were matched using absolute parcel-wise Pearson correlations, allowing for sign reversal and differences in ordering. Correspondence between the four-dimensional loading subspaces was additionally quantified by principal angles between orthonormal loading bases.

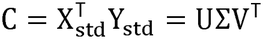

The dimensional spectrum of the symmetric decomposition was summarized by the squared singular values, normalized by their sum. Cumulative shared cross-covariance structure was calculated from these normalized values across successive components.

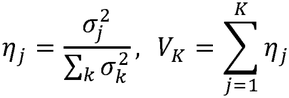

### Participant-level reproducibility of PLS components

To evaluate the influence of participant composition, we repeated the PLS decomposition under leave-one-participant-out, independent split-half and participant-bootstrap resampling. Functional-connectivity matrices were first averaged across runs within each participant. Leave-one-participant-out analysis recomputed the group-level matrices and PLS loadings after excluding each of the 17 participants in turn. Split-half analysis used 1,000 randomly generated non-overlapping partitions of the participants into groups of eight and nine, with the decomposition fitted independently in each half. Participant bootstrap resampling generated 1,000 samples of 17 participants with replacement and refitted the decomposition for each sample.

Resampled loading vectors were matched to the corresponding full-sample components by absolute spatial correlation, accounting for component ordering and sign indeterminacy. Reproducibility was summarized by the correlations of matched loading maps, with split-half correspondence evaluated between the independent subsets.

Bootstrap loading estimates were retained for the parcel-level stability analysis.

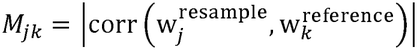

### Permutation-based thresholding of regional PLS loadings

Regional loading contributions were evaluated using a non-parametric permutation procedure with 1,000 permutations separately for auditory and visual loading matrices. For each permutation, rows of the response matrix were randomly shuffled, the four-component PLS model was refit, and the resulting response loadings were stored. Let 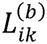 denote the loading for parcel *i*, LV *k*, and permutation *b*. Element-wise two-sided null limits were defined from the 2.5th and 97.5th percentiles,

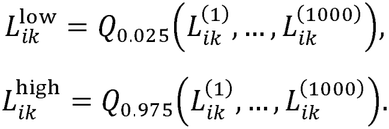

The observed loading 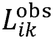 was retained if it fell outside this interval and was otherwise otherwise set to zero,

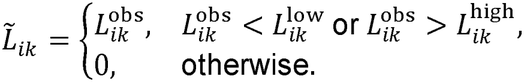

This corresponds to an elementwise, two-sided permutation-derived threshold of *α* = 0.05, uncorrected for multiple comparisons. The native component orientation returned by the estimator was retained in each permutation; no post hoc sign alignment was applied. Independent fixed random seeds were used for the auditory and visual permutation procedures.

### Participant-bootstrap stability of regional loadings

For each parcel, modality and latent variable, the 2.5th and 97.5th percentiles of participant-bootstrap loadings defined a 95% confidence interval. Parcels whose intervals excluded zero were classified as bootstrap-stable. To compare this stability criterion with the permutation-based loading threshold, we calculated the proportion of permutation-selected parcels recovered in the corresponding bootstrap-stable set and their loading-sign agreement. The proportions of selected parcels were also compared across modalities and components.

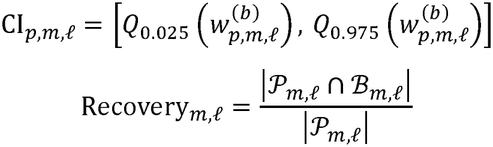

**K-means analysis of the LV1-LV2 loading space**

To test whether the geometry of the two dominant auditory loading dimensions yielded two separable parcel populations independently of the permutation labels, K-means clustering was performed in the auditory LV1-LV2 raw loading space. Parcel *i* was represented by

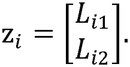

Only parcels with a non-zero permutation-thresholded loading in LV1, LV2, or both were entered into K-means. Two clusters were fitted with 50 initializations and random seed 42. The objective was to minimize within-cluster squared Euclidean distance,

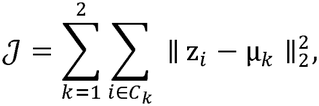

where *C*_*k*_ is cluster *k* and µ_*k*_ is its centroid.

Independently of K-means, parcels were categorized from the permutation results as LV1-only, LV2-only, significant in both LVs, or nonsignificant in both. Parcels significant in both LVs were displayed in loading space but excluded from the binary agreement analysis because they did not provide an exclusive LV1-versus-LV2 assignment.

Because K-means labels are arbitrary, cluster identities were relabeled where necessary to maximize correspondence with the LV1-only and LV2-only significance categories. Agreement was defined as

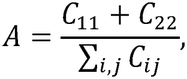

where C is the 2 × 2 contingency matrix after optimal label correspondence. Adjusted Rand index and normalized mutual information were additionally calculated as label-invariant measures of partition similarity.

Statistical significance of the observed agreement was assessed using 5,000 label permutations. At permutation *b*, the exclusive LV1/LV2 significance labels were shuffled while K-means memberships were held fixed. Because the two cluster labels remain exchangeable under the null, both the identity and swapped label correspondences were evaluated, and the larger agreement was retained. The probability of permutation

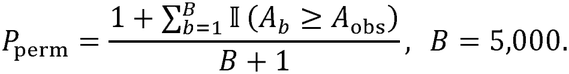

### Correspondence between FC clusters and PLS-significant parcels

The relationship between hierarchical FC clusters and the dominant PLS dimensions was quantified by measuring the fraction of significant LV parcels contained within each FC cluster. For FC cluster *C*_*j*_ and the set *S*_*k*_ of permutation-significant parcels for LV *k*,

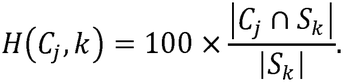

Thus, the denominator is the total number of significant parcels for the corresponding LV. Each heat-map cell represents the percentage of LV’s significant regional contributors contained in a given FC cluster.

### Cole-Anticevic brain-wide network analysis

Large-scale network affiliation of PLS loadings was characterized using the Cole-Anticevic Brain-wide Network Partition (CAB-NP)^25^, which assigns HCP-MMP cortical parcels to 12 large-scale functional networks: Visual1, Visual2, Auditory, Somatomotor, Dorsal-attention, Cingulo-Opercular, Frontoparietal, Language, Default, Posterior-Multimodal, Ventral-Multimodal and Orbito-Affective.

Because the atlas used in the present analysis combined homologous left- and right-hemisphere territories into 180 symmetric parcel identities, a conservative bilateral network-assignment rule was applied. For each symmetric parcel, the CAB-NP assignments of the corresponding left- and right-hemisphere HCP-MMP areas were compared. A symmetric parcel was assigned to a CAB-NP network only when both hemispheric homologues had the same network label. Of 180 parcels, 167 had concordant left-right CAB-NP assignments and were retained, whereas 13 discordant parcels were excluded from this network-level analysis. All 12 CAB-NP networks remained represented.

For network *g*, modality *m*, and LV *k*, the network-level loading profile was

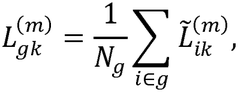

where *N*_*g*_ is the total number of retained parcels assigned to network *g* and 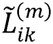 is the signed permutation-thresholded loading. Nonsignificant parcels therefore entered the network average as zeros rather than being removed. The network statistics consequently reflect both the magnitude of significant loadings and how broadly those loadings are distributed across parcels within the network.

### Latent-variable graph construction

To characterize the connectivity architecture associated with each PLS dimension, a separate undirected graph was constructed for each modality, and LV. Nodes were restricted to parcels with non-zero permutation-thresholded loading on the corresponding LV,

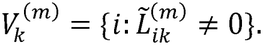

For each pair of retained nodes, the candidate edge weight was taken from the corresponding group-average FC matrix,

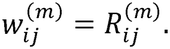

An adaptive threshold was defined separately for each modality-LV graph,

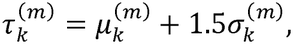

where 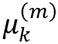 and 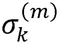 are the mean and standard deviation of all candidate FC weights connecting the retained nodes. An edge was included when

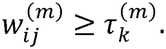

Thresholding was applied to the original signed correlation coefficients rather than their absolute values; therefore, the final graphs retained strongly positive and negative functional connections. Nodes without any edges above the threshold were removed from analysis.

### Graph degree, edge overlap and core subnetworks

For node *i*, degree was

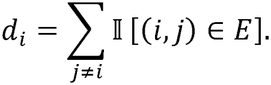

Degree distributions were compared between auditory and visual LV1 graphs, auditory and visual LV2 graphs, and the auditory LV3 and visual LV4 graphs using two sample Kolmogorov-Smirnov tests. The test statistics were

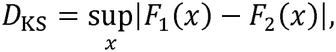

where *F*_l_ and *F*_2_ are the corresponding empirical cumulative degree distributions. Cross-modal edge conservation was summarized with an overlap coefficient,

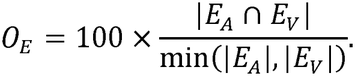

This measure quantifies the percentage of edges in the smaller graph that are also present in the larger graph. It is distinct from the Jaccard index.

For chord-diagram visualization, core subnetworks were defined as the top 15% of nodes ranked by degree. If *V* denotes the node set,

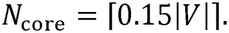

For LV1 and LV2, a shared auditory-visual graph was defined from the intersection of the two modality-specific edge sets. For edges present in both modalities, the visualization weight was the mean of their auditory and visual FC weights,

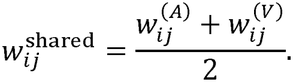

The highest-degree induced subgraph was then extracted. Auditory LV3 and visual LV4 cores were obtained separately from their respective graphs. Absolute edge magnitudes were used to scale chords for visualization only; graph selection itself was based on the signed positive-FC threshold described above.

### Temporal expression of the connectivity latent variables

To link static connectivity structure to time-resolved BOLD dynamics, each run was projected onto the modality-specific raw, unthresholded PLS loading matrix. Let

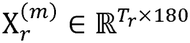

be the parcel-level BOLD matrix for run *r* and modality *m*, and let

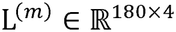

contain the corresponding four PLS loading vectors. Temporal latent-variable expressions were computed as

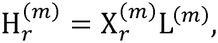

yielding four latent time series (tLV1-tLV4). For component *k*,

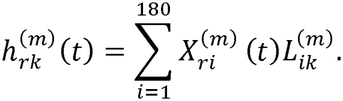

Permutation thresholding was therefore used to interpret spatial contributions but was not imposed on the loading matrix before temporal projection.

For the dynamical analyses, each tLV was standardized within each run,

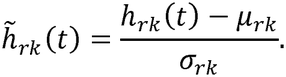

### Spectral characterization of tLV dynamics

Frequency content of the raw tLVs was characterized using the discrete Fourier transform. Runs were truncated to a common length where necessary. For tLV *k*,

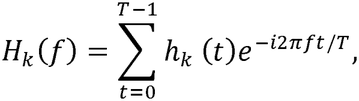

and spectral power was defined as

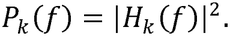

Only non-negative frequencies were retained, and spectra were averaged across runs. With TR = 2 s, the sampling frequency was 0.5 Hz, and the Nyquist frequency was 0.25 Hz. No additional spectral normalization was applied in the final figure-generation analysis.

### Vector-autoregressive modelling of latent dynamics

The main dynamical analysis focused on the standardized tLV1-tLV2 state vector,

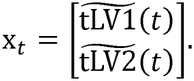

Autoregressive orders *p* = 1,2,3,4 were examined during exploratory model development. A second-order model was retained as a parsimonious final representation incorporating the previous two sampled states. The intrinsic VAR(2) model was

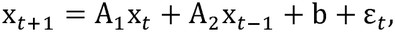

where A_1_ and A_2_ are 2 × 2 lag matrices, b is an intercept vector, and ε_*t*_ is residual error. Parameters were estimated using multivariate ridge regression with *α* = 1. Writing the lagged design matrix as Z and the stacked coefficient matrix as Θ = [A_1_ A_2_], parameter estimation minimizes

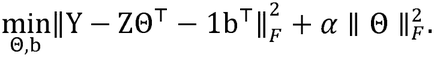

Predictive performance was assessed using five-fold blocked cross-validation. Lagged observations were divided into contiguous test blocks, and each held-out block was predicted from a model fitted to the remaining blocks. This procedure is described as blocked rather than forward-chaining cross-validation because training observations could occur on both temporal sides of a held-out block. For the two-dimensional output, scikit-learn’s uniform-average multi-output coefficient of determination was used,

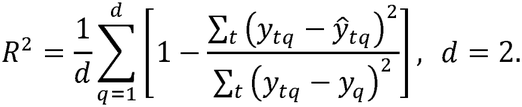

### Target events as an external dynamical input

To test whether target events explained state evolution beyond intrinsic autoregressive history, a binary target input was constructed from the event timing files. Each target onset was converted to a BOLD sample index using the 2-s TR and shifted by one TR. The one-TR delay was retained after exploratory comparison of candidate temporal shifts and was used consistently in the final controlled model. This discrete shift provides a simple alignment with delayed BOLD expression and was not intended to substitute for explicit haemodynamic-response-function convolution.

The controlled VAR(2) model was

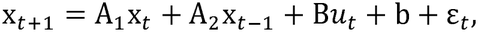

where *u*_*t*_ ∈ {0,1} is the shifted target indicator and B ∈ ℝ^2 × 1^ represents the target-associated input. The same ridge regularization and blocked cross-validation scheme were used for models with and without the external input. Incremental predictive contribution was summarized as

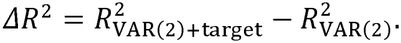

### Dynamical stability and fixed points

Stability of each fitted VAR(2) system was assessed using the companion matrix

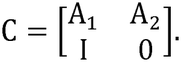

The fitted dynamics were classified as stable when the spectral radius satisfied

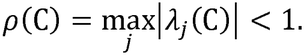

For a constant external input *u*, a fixed point x*(*u*) satisfies

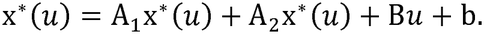

Thus,

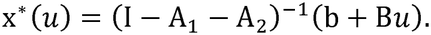

The baseline fixed point is

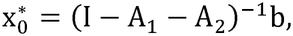

and the target-conditioned fixed point is

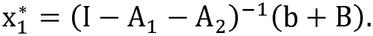

The corresponding model-implied target displacement is

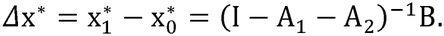

Fixed points were estimated at the run level and summarized across runs.

### Baseline latent-state landscape

To visualize empirical occupancy of the tLV1-tLV2 state space, within-run standardized states were pooled across runs after excluding samples marked by the shifted target regressor. The resulting target-free observations were used to estimate a two-dimensional Gaussian kernel density,

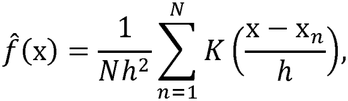

where x_*n*_ is an observed state and *h* is the KDE bandwidth. Density was evaluated on a 220 220 grid with 20% spatial padding. Contours corresponding to 50%, 75%, 90% and 95% cumulative probability mass were used for visualization. The mean run-wise baseline fixed point was superimposed on this empirical density landscape. Thus, the displayed density is an empirical KDE of target-free state occupancy; the VAR model contributes the fixed-point estimate but does not generate the density surface itself.

### Gaussian-mixture characterization of latent configurations

To provide a coarse-grained description of preferential occupancy within the continuous latent space, Gaussian mixture models (GMMs) were fitted to the raw four-dimensional tLV vectors pooled across all 101 runs. With 170 samples per retained run, this yielded 17,170 four-dimensional observations.

For *K* components,

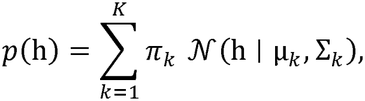

where π_*k*_ is the mixture weight, µ_*k*_ the four-dimensional centroid and Σ_*k*_ the full covariance matrix of component *k*.

Candidate solutions with *K* = 1, …, 5 were inspected during exploratory development using log likelihood and the Akaike and Bayesian information criteria,

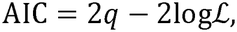

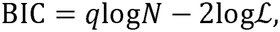

where *q* is the number of fitted parameters and is the number of observations. A three-component solution was retained because it provided a parsimonious, interpretable coarse-graining of positive, central and negative configurations along the dominant latent dimensions. The final GMM used full covariance matrices, 20 random initializations and random seed 42. Posterior membership probabilities were

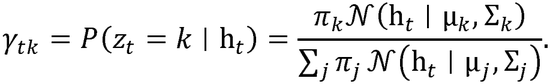

Hard labels were assigned by maximum posterior probability,

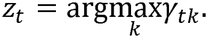

Because GMM labels are arbitrary, components were ordered according to the sum of their tLV1 and tLV2 centroid coordinates,

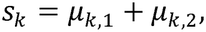

with descending values labelled positive, central and negative, respectively. A temporal sample was additionally classified as high confidence when

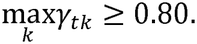

This confidence threshold was descriptive and did not alter the fitted mixture model.

### Latent-configuration occupancy and transition probabilities

Overall occupancy of configuration *k* was 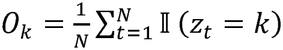.

Transitions were counted only between consecutive samples within the same run, preventing artificial transitions across run boundaries. The count of transitions from configuration *i* to configuration *j* was

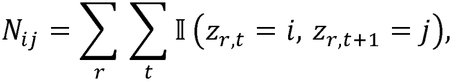

and the row-normalized transition probability was

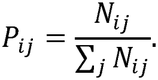

Diagonal terms *P*_*ii*_ therefore quantify persistence, whereas off-diagonal terms describe movement between regions of preferential occupancy.

### UMAP visualization of four-dimensional latent dynamics

The full raw four-dimensional tLV representation was visualized using Uniform Manifold Approximation and Projection^58^. The final embedding used two output dimensions, 15 nearest neighbours, min dist *= 0*.*1*, and random seed 42. The embedding was fitted to all available four-dimensional tLV observations; any subsequent point subsampling was used only to reduce overplotting in the displayed scatterplots.

UMAP constructs a fuzzy neighbourhood representation in the original space and optimizes a low-dimensional embedding that preserves those neighbourhood relations. Its low-dimensional similarity can be written generically as

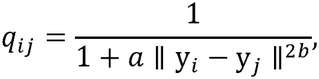

with the embedding chosen to minimize fuzzy-set cross-entropy between high- and low-dimensional neighbourhood memberships. UMAP was used as a descriptive visualization and not as an inferential test of dimensionality or discrete state number.

To examine modality-related organization independently of the two dominant latent dimensions, the cross-modal PLS decomposition was repeated with five components. Each auditory and visual run was projected onto its corresponding five-component loading matrix, and tLV3–tLV5 were retained for a separate UMAP analysis. The three-dimensional input was embedded into two dimensions using the same UMAP parameters as the original four-dimensional analysis (15 nearest neighbors, minimum distance = 0.1, random seed = 42). Auditory and visual observations were projected jointly, and their distributions were compared descriptively within the resulting embedding.

### Posterior-weighted configuration density in UMAP space

To visualize the relationship between the continuous manifold and GMM configurations without relying only on hard labels, posterior probabilities were used as continuous weights in two-dimensional kernel-density estimates. For UMAP coordinate *y* and configuration *k*,

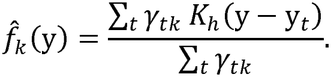

Thus, each sample contributed to each configuration surface in proportion to its posterior membership probability. The density fields were Gaussian-smoothed for visualization, and their maxima were used to identify density peaks in the UMAP plane. The GMM itself remained fitted in the original four-dimensional tLV space; UMAP coordinates were used only for visualization of that fitted structure.

### Modality-dependent covariance structure in latent space

To compare the organization of auditory and visual temporal trajectories, raw tLV observations from both modalities were first pooled and each LV was standardized globally across all samples,

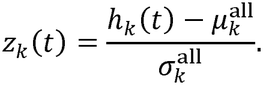

Auditory and visual covariance structure was then examined separately in the tLV1-tLV2 and tLV3-tLV4 planes. For modality *m*,

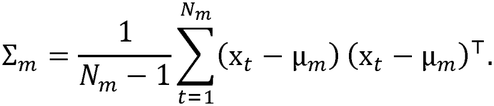

Principal covariance directions were obtained by eigen decomposition,

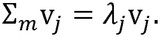

Ellipses were oriented by the eigenvectors v and drawn with full axis lengths 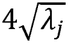, corresponding to semi-axes of 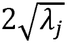. This analysis was descriptive and assessed the orientation and dispersion of modality-specific trajectories within a common standardized latent space.

### Target-associated displacement in the tLV1-tLV2 landscape

The controlled VAR model was used to visualize the direction of target-associated displacement relative to the baseline fixed point. States were recentered as

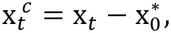

such that the baseline fixed point lay at the origin. The model-implied target displacement vector

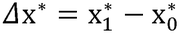

was displayed together with target-associated observations. The representative single-run display was used to illustrate the geometry of the fitted model; quantitative cross-validated performance, stability and fixed-point summaries were evaluated across runs.

### Overlap-adjusted finite impulse response analysis

Target- and standard-associated BOLD trajectories were estimated using a joint finite impulse response (FIR) model. Actual stimulus onset times were reconstructed from the experiment event records and checked against run identities, event counts and acquisition timing. Each eligible run contained 25 targets and 100 standard onsets. Of the 101 retained functional runs, 91 had verified event records suitable for the FIR analyses (48 auditory and 43 visual runs, representing all 17 participants).

Because successive stimuli were separated by only 2–3 s, event-related responses overlapped in time. We therefore fitted separate target and standard FIR regressors simultaneously within each run, allowing all verified events to contribute to the modeled BOLD signal. The basis comprised causal piecewise-linear functions evaluated at the exact event onsets, with knots every 2 s from 0 to 24 s after stimulus onset. The design matrix also included an intercept and linear run trend. Ordinary least squares was used to estimate target and standard coefficients for the within-run standardized tLV1 and tLV2 time series; models were required to have full-column rank, a scaled-design condition number below 10□ and sufficient residual degrees of freedom.

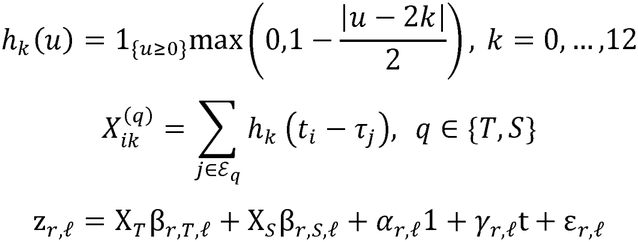

The resulting target and standard coefficient sequences were interpreted as their modeled event-associated BOLD contributions at each post-stimulus lag. For each latent dimension, the target–standard trajectory was obtained by subtracting the standard FIR coefficients from the target FIR coefficients at the same lag. These contrasts accounted for the modeled temporal overlap between neighboring stimuli without requiring a canonical hemodynamic response shape.

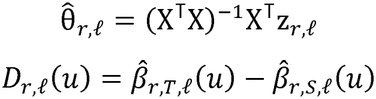

Participant-level target–standard contrasts

FIR coefficients were first averaged across eligible runs within each participant and modality, and participant averages were then used for group summaries and inference. The target–standard difference at the 4-s lag was examined separately for auditory and visual tLV1. The late contrast was defined as the change in the target–standard difference between 4 and 6 s, [(target_6_ − standard_6_) − (target_4_ − standard_4_)], and calculated for both dominant latent dimensions. Target-only and standard-only 4–6-s changes were also retained to distinguish their contributions to the late contrast.

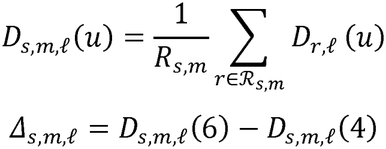

Two-sided participant-level sign-flip tests assessed the mean contrasts, with exact enumeration of sign assignments for analyses based on the 17 paired participants. The 4-s tLV1 comparisons were Holm-adjusted across the two modalities. For the late response, auditory–visual differences were tested within participants using the paired difference between their 4–6-s contrasts. Participant bootstrap resampling provided percentile confidence intervals for the group trajectories and contrast estimates. The 4-s and subsequent 4–6-s analyses were identified from inspection of the estimated event-related trajectories and treated as exploratory.

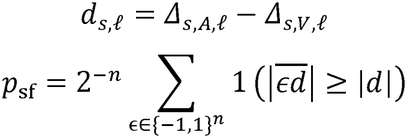

Parcel contributions to early and late latent trajectories

Regional contributions were derived from the same 24-s FIR design fitted separately to each parcel-wise BOLD time series. At each lag, the estimated parcel response was multiplied by its unthresholded PLS loading for the corresponding modality and latent variable, then divided by the standard deviation of that run’s projected tLV. Summing these weighted parcel contributions recovered the estimated tLV FIR coefficient. The 4-s parcel contribution was calculated from the target–standard coefficient difference, whereas the late contribution was the difference between the 6-s and 4-s target– standard contributions.

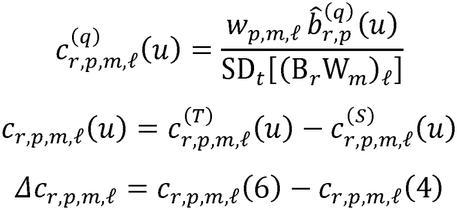

Parcel estimates were averaged within participants and then across participants for each modality. For the 4-s tLV1 maps, the 15 strongest positive contributors were selected independently in auditory and visual runs and their union displayed using a common parcel mask and color scale. To examine the late response across both dominant dimensions, parcels were ranked by the absolute magnitude of their auditory– visual-pooled 4–6-s contributions separately for tLV1 and tLV2. The union of the 15 highest-ranked parcels from each dimension contained 27 unique parcels. For the joint anatomical display, each parcel was assigned to the dimension with the larger absolute pooled contribution when it appeared in both selections, and assignments were held fixed across modalities.

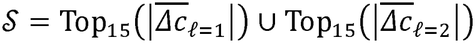

For statistical comparisons, the signed contributions of selected parcels were summed separately along tLV1 and tLV2 for each participant and modality. The two-dimensional auditory–visual difference was tested using the Euclidean norm of the participant-mean paired contribution vector, with an exact two-sided sign-flip distribution formed from all 2^1^□ sign assignments. Parallel analyses of the 15 and 20 largest pooled two-dimensional contributors provided selection-size comparisons, with Holm correction across the three joint tests. Differences in vector magnitude were tested separately from differences in the signed contribution vector. The 15 strongest positive and negative parcel contributors were also compared between modalities for each dimension using participant-level sign-flip tests, with Holm correction across the four signed-contributor comparisons.

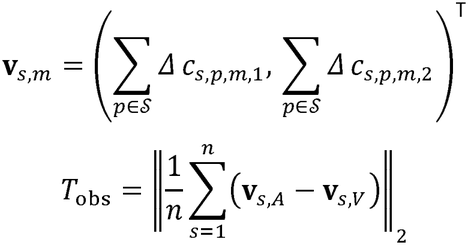

### Sensitivity of late event-related estimates

Sensitivity of the 4–6-s contrasts to response-model duration was assessed by repeating the joint FIR analysis with supports of 16, 20, 24 and 28 s, using the same eligible runs throughout. A conventional double-gamma hemodynamic response model with its temporal derivative, fitted separately for actual target and standard onsets, provided an additional comparison. The 24-s FIR solution was used for the parcel-wise contribution analyses and the main spatial visualizations.

