## Supplementary Materials for "Sensory Change Detection Unfolds Within a Shared Low-Dimensional Brain-State Landscape"

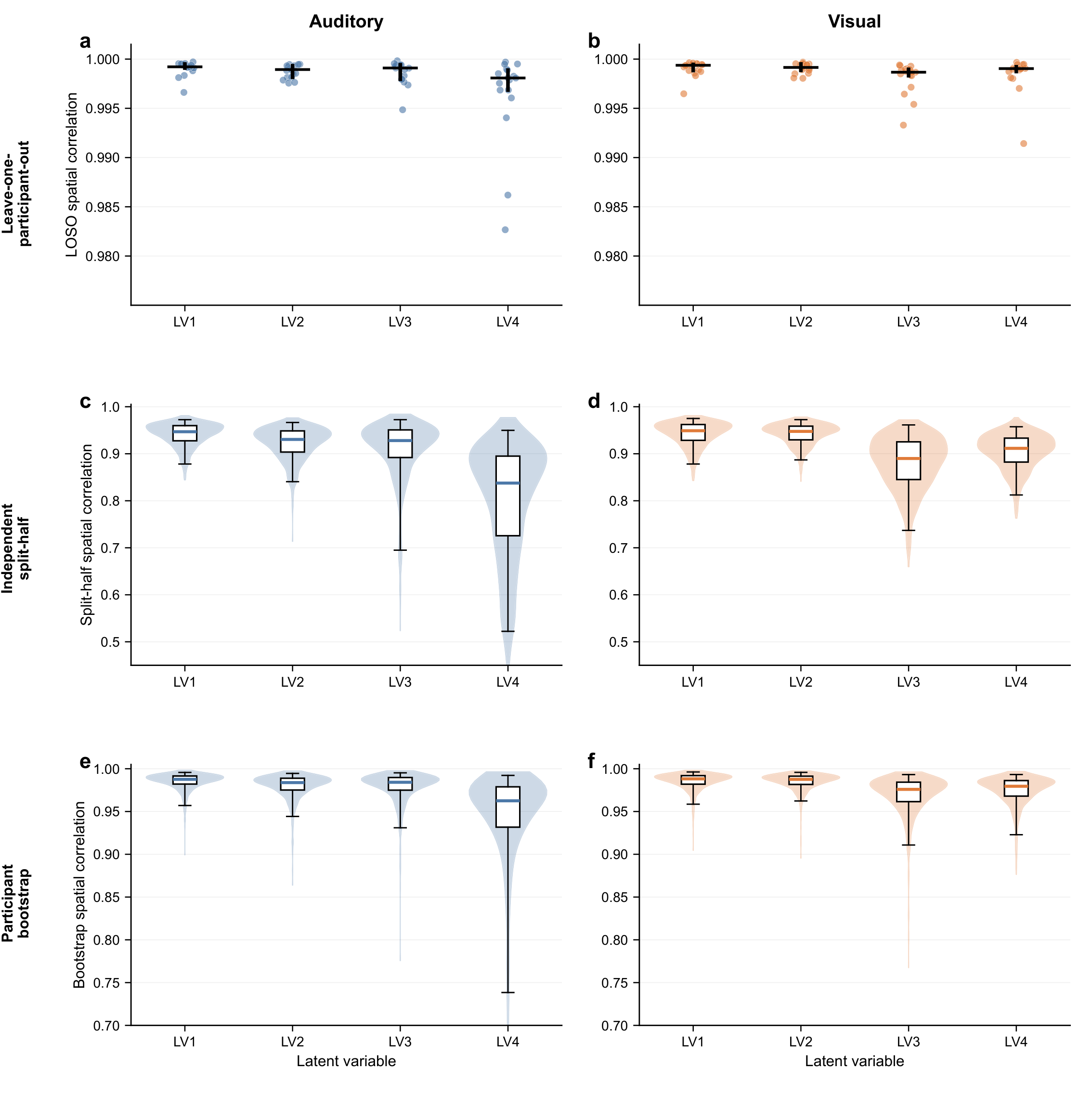

**Supplementary Fig. 1 | Participant-level reproducibility of PLS loading maps.** Auditory (a,c,e) and visual (b,d,f) loading maps were compared across leave-one-participant-out samples (a,b), 1,000 independent split-half partitions (c,d) and 1,000 participant-bootstrap resamples (e,f). Spatial correlations quantify loading-map correspondence.

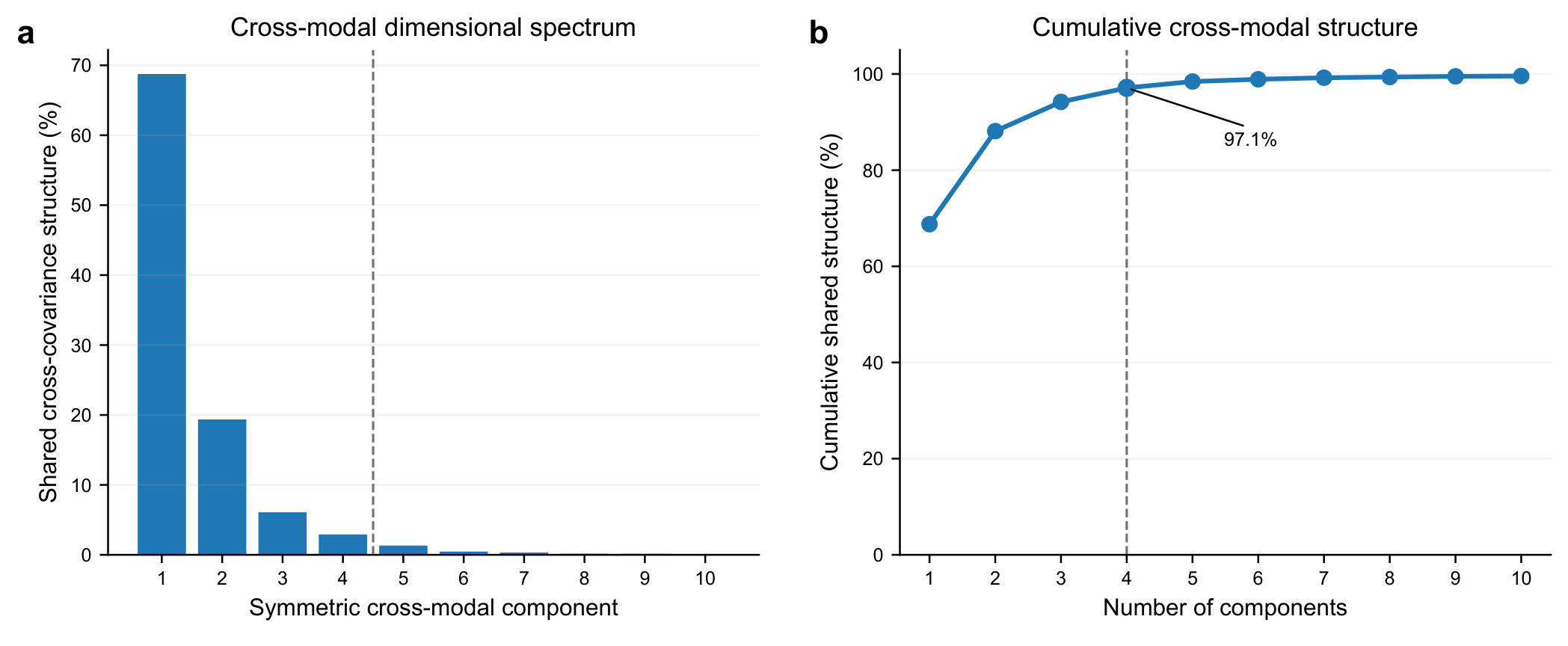

**Supplementary Fig. 2 | Dimensionality of the symmetric cross-modal decomposition.** a, Proportion of shared cross-covariance structure captured by successive symmetric components. b, Cumulative structure, with 97.1% captured by the first four components (dashed line).

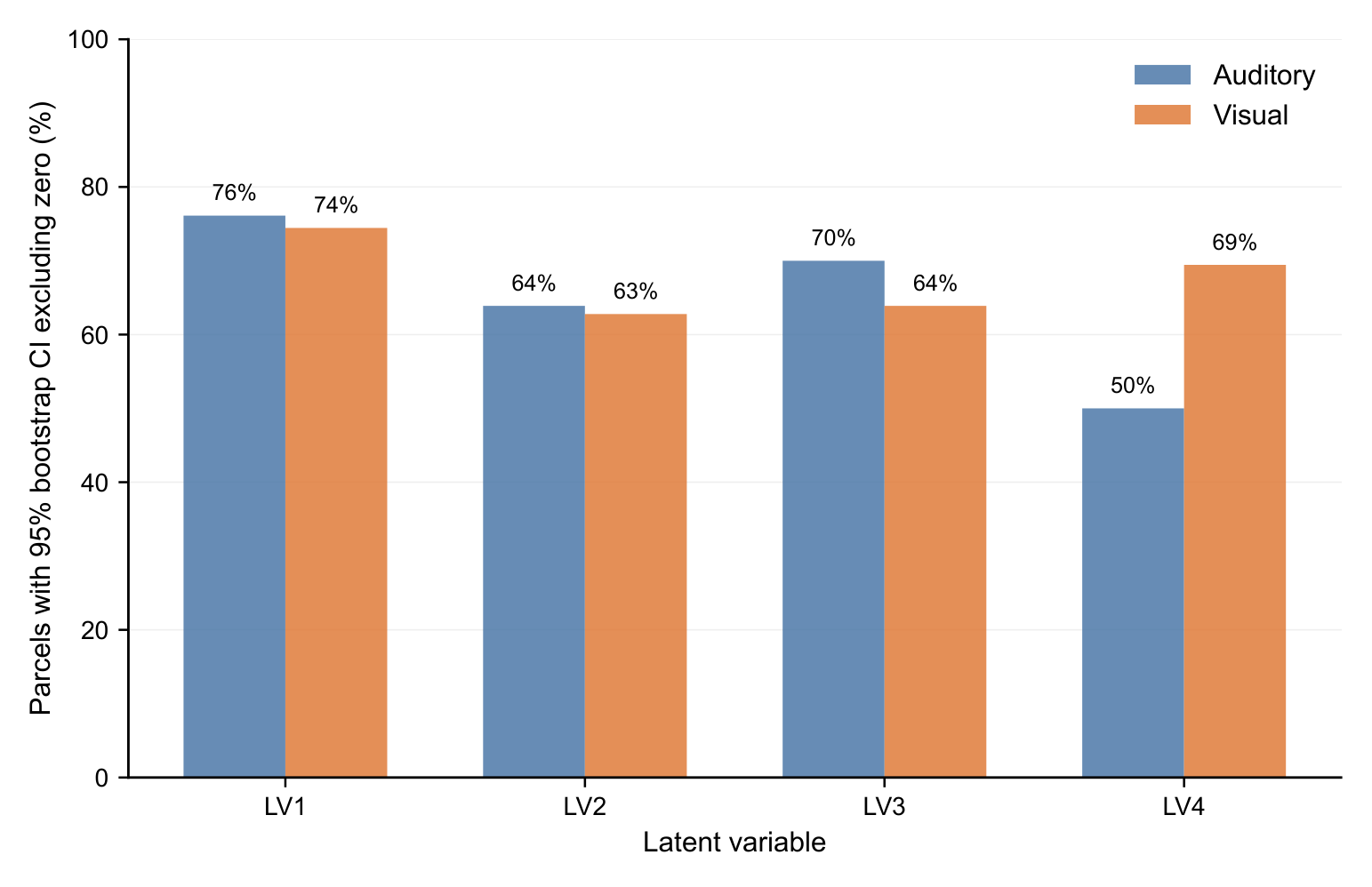

**Supplementary Fig. 3 | Stability of parcel-wise PLS loadings.** Percentage of cortical parcels with participant-bootstrap 95% confidence intervals excluding zero, shown for LV1–LV4 in auditory and visual conditions.

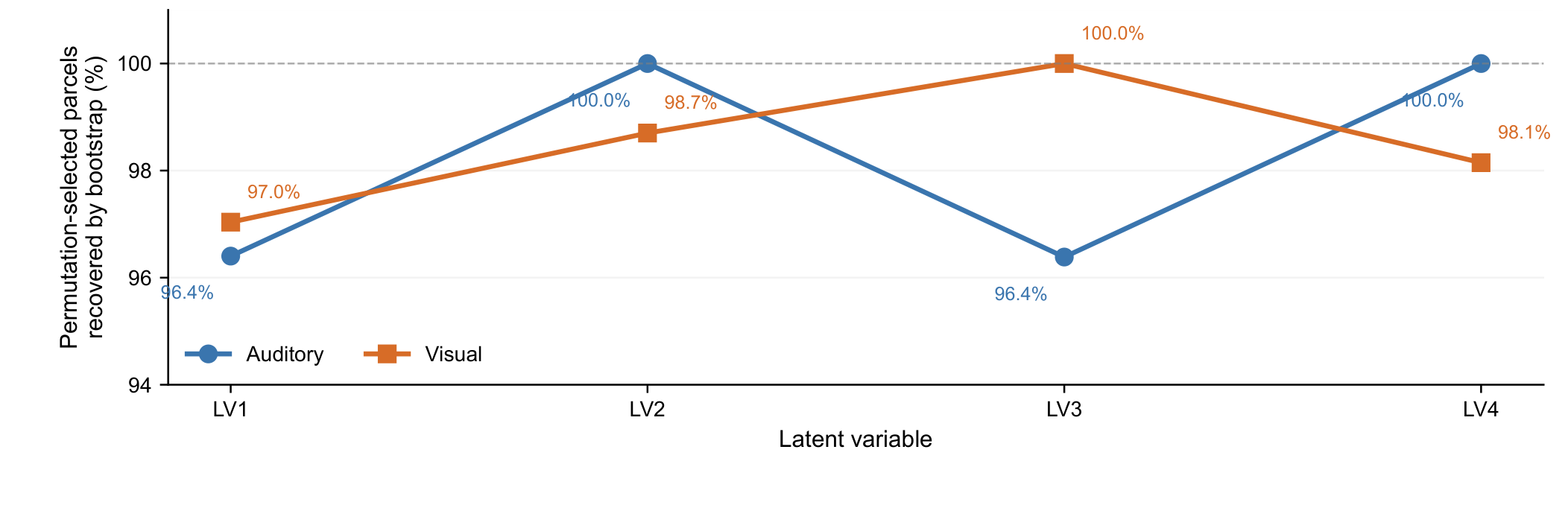

**Supplementary Fig. 4 | Recovery of permutation-selected parcels by participant bootstrap.** Percentage of permutation-selected parcels also identified as bootstrap-stable in each modality and latent variable.

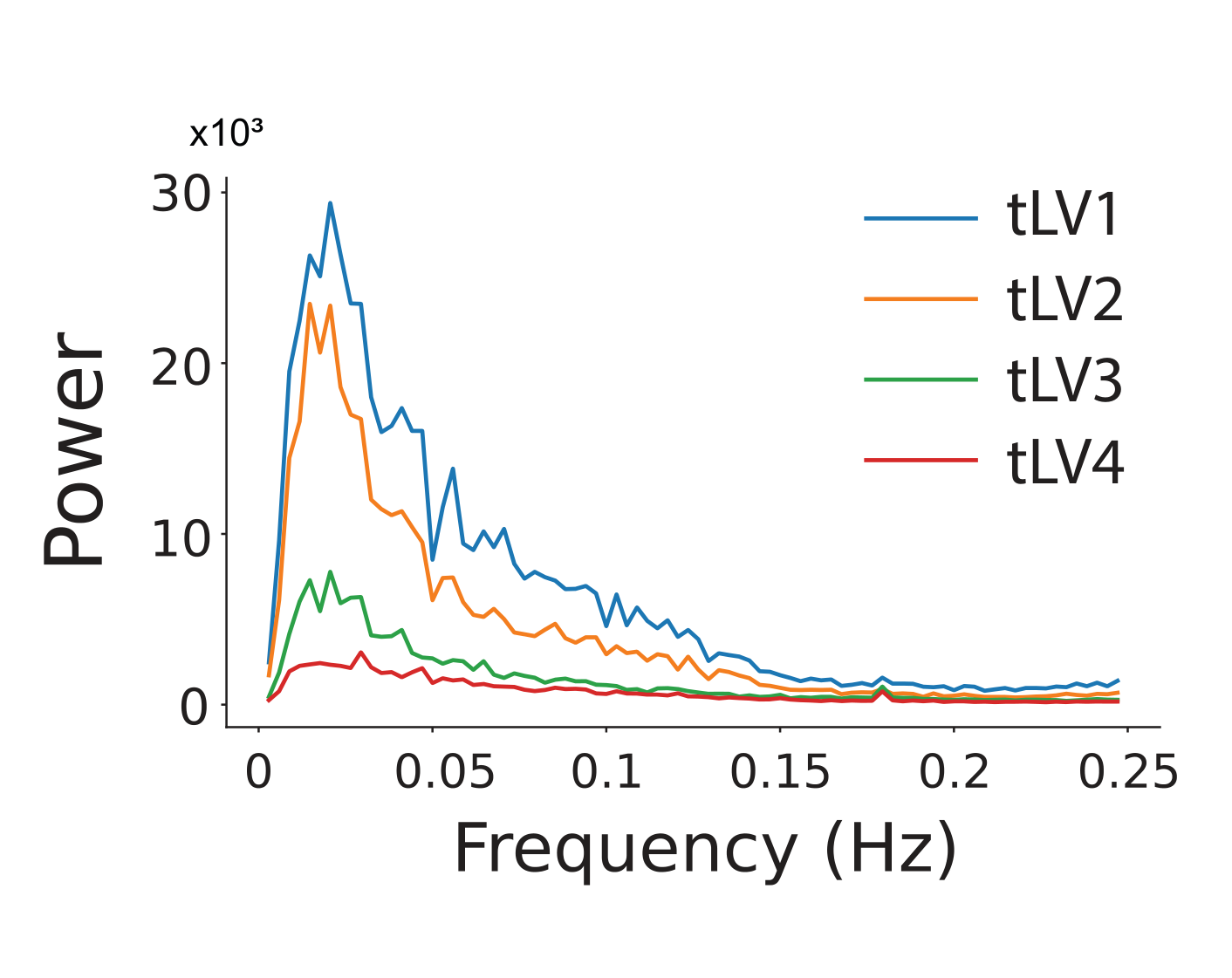

**Supplementary Fig. 5 | Spectral power of temporal latent variables.** Mean power spectra for tLV1–tLV4 across runs, plotted over the sampled frequency range (TR = 2 s).

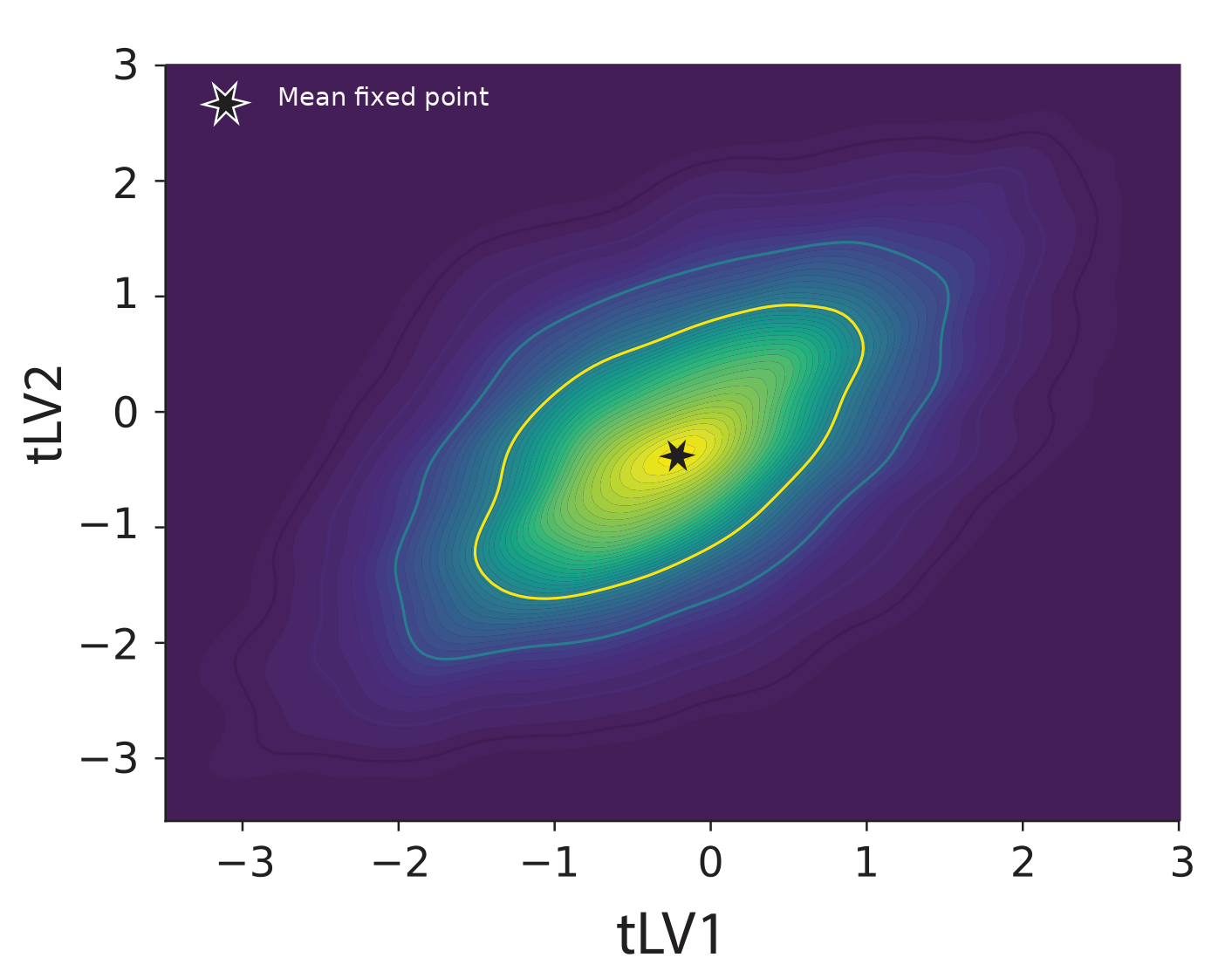

**Supplementary Fig. 6 | Empirical occupancy in the dominant temporal latent space.** Density of pooled target-free observations in the standardized tLV1–tLV2 plane. Contours show regions of higher occupancy; the star marks the mean fitted baseline fixed point.

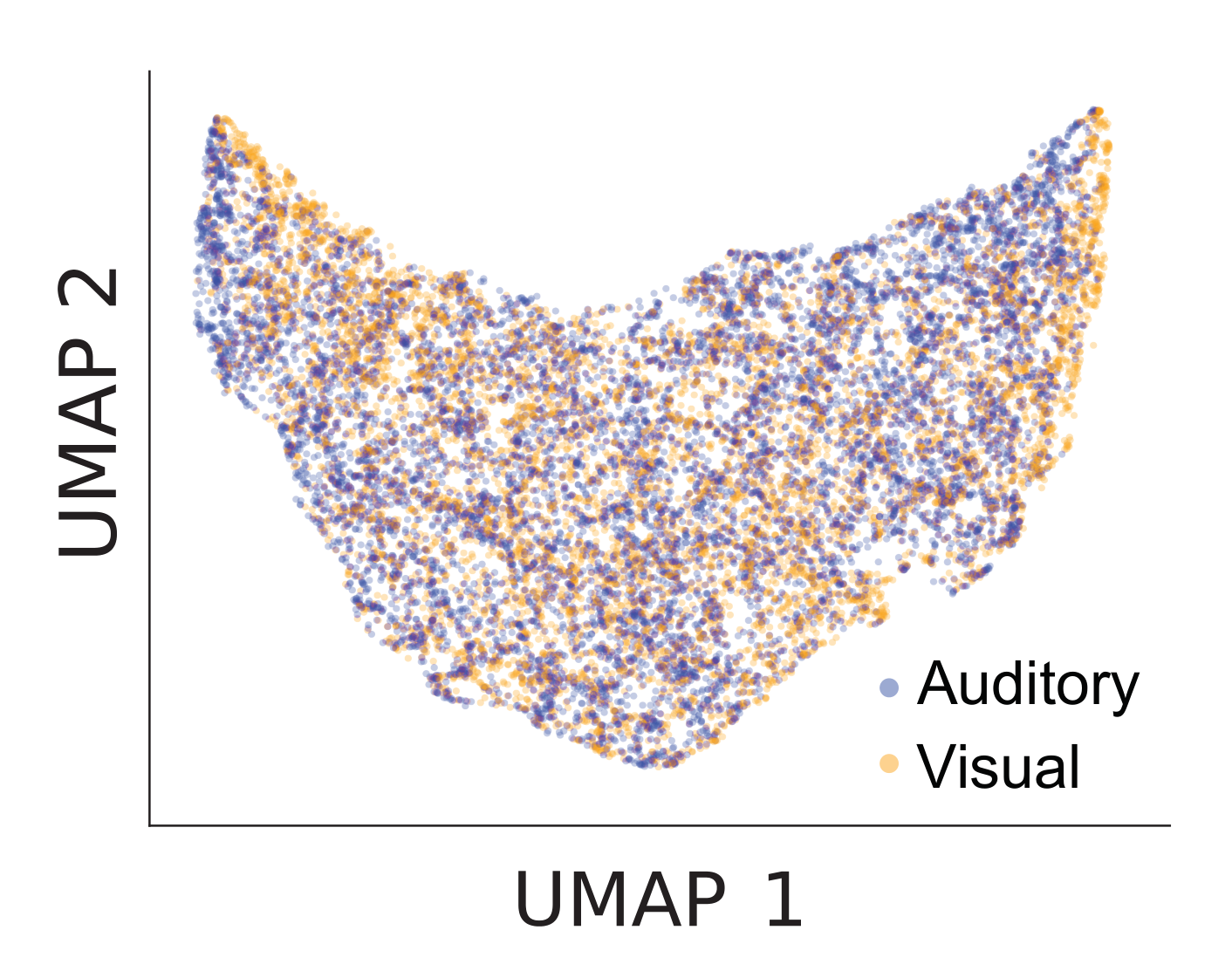

**Supplementary Fig. 7 | Auditory and visual distributions in the four-dimensional tLV embedding.** Two-dimensional UMAP representation of pooled tLV1–tLV4 observations, coloured by task modality. Auditory and visual samples occupy largely overlapping regions of the embedding.
